# The role of sigma-1 receptor in organization of endoplasmic reticulum signaling microdomains

**DOI:** 10.1101/2020.12.04.411553

**Authors:** Vladimir Zhemkov, Jonathon A. Ditlev, Wan-Ru Lee, Jen Liou, Michael K. Rosen, Ilya Bezprozvanny

## Abstract

Sigma 1 receptor (S1R) is a 223 amino acid-long transmembrane endoplasmic reticulum (ER) protein. S1R modulates activity of multiple effector proteins but its signaling functions are poorly understood. We here test the hypothesis that biological activity of S1R in cells can be explained by its ability to interact with cholesterol and to form cholesterol-enriched microdomains in the ER. Using reduced reconstitution systems, we demonstrate direct effects of cholesterol on S1R clustering. We identify a novel cholesterol-binding motif in the transmembrane region of S1R and demonstrate its importance for S1R clustering. We demonstrate that S1R-induced membrane microdomains have increased local membrane thickness. Increased local cholesterol concentration and membrane thickness in these domains can modulate signaling of inositol-requiring enzyme 1α (IRE1α) in the ER. Further, S1R agonists cause reduction in S1R clusters, suggesting that biological activity of S1R agonists is linked to remodeling of ER membrane microdomains.

## INTRODUCTION

Cholesterol is an essential component of cellular membranes and levels of cholesterol in cells are tightly controlled (Brown & Goldstein, 1999; Goldstein & Brown, 2015; Radhakrishnan, Goldstein, McDonald, & Brown, 2008). Plasma membranes are highly enriched in cholesterol and other sterols and contain 30-40% cholesterol and 10-30% sphingolipids based on molar amounts (van Meer, Voelker, & Feigenson, 2008). It is widely accepted that cholesterol is not distributed homogeneously in the plasma membrane and instead forms lipid rafts, or cholesterol-enriched microdomains, that play important signaling roles in cells (Levental, Levental, & Heberle, 2020). Although cholesterol is produced in the endoplasmic reticulum (ER), concentration of cholesterol in ER membranes is much lower than in plasma membranes (van Meer et al., 2008). Level of ER cholesterol is maintained by on-and-off switch of the Scap-SREBP pathway with half-maximal response of SREBP-2 at about 5% molar cholesterol in the ER (Radhakrishnan et al., 2008). There is limited information about spatial distribution of cholesterol in the ER membrane, but recent studies suggested existence of cholesterol-enriched microdomains in the ER membrane (Area-Gomez et al., 2012; Hayashi & Fujimoto, 2010; King, Sengupta, Seo, & Lippincott-Schwartz, 2020; Montesinos et al., 2020). One of the most studied examples of such microdomains are mitochondria-associated membranes (MAMs), which are sites on the endoplasmic reticulum (ER) membrane in immediate proximity to the mitochondrial outer membrane (Csordas, Weaver, & Hajnoczky, 2018; Prinz, Toulmay, & Balla, 2020). MAMs have been shown to play an important role in a variety of cellular functions, such as ER to mitochondria Ca^2+^ transfer (Csordas et al., 2018; Hajnoczky, Csordas, & Yi, 2002), ATP production (Hajnoczky et al., 2002), lipid metabolism (Csordas et al., 2018; Vance, 2014) and autophagy (Garofalo et al., 2016). Not surprisingly, MAM dysregulation is observed in numerous pathophysiological conditions, including neurodegenerative diseases (Schon & Area-Gomez, 2013), cancers (Morciano et al., 2018) and lysosomal disorders (Annunziata, Sano, & d’Azzo, 2018; Sano et al., 2009).

Sigma 1 receptor (S1R) is a 223 amino acid-long single-pass transmembrane ER protein (Hayashi, 2019; D. A. Ryskamp, Korban, Zhemkov, Kraskovskaya, & Bezprozvanny, 2019; Schmidt & Kruse, 2019) that has a short cytoplasmic tail and a large luminal ligand binding domain (Mavylutov, Chen, Guo, & Yang, 2018; Schmidt et al., 2016). It has been suggested that S1R acts as a molecular chaperone that can stabilize a native conformation of multiple client proteins in stress conditions (Hayashi, 2019; Nguyen, Lucke-Wold, Mookerjee, Kaushal, & Matsumoto, 2017; T. P. Su, Hayashi, Maurice, Buch, & Ruoho, 2010). Recent studies also suggested that S1R can act as an RNA binding protein (P. T. Lee et al., 2020). S1R is highly enriched in liver and expressed in the nervous system. Analysis of S1R knockout mice revealed a number of nervous system abnormalities (Couly et al., 2020), suggesting that S1R plays an important role in neurons. Mutations in S1R lead to a juvenile form of ALS (Al-Saif, Al-Mohanna, & Bohlega, 2011) and distal hereditary motor neuropathy (Almendra, Laranjeira, Fernandez-Marmiesse, & Negrao, 2018; Gregianin et al., 2016; Horga et al., 2016; X. Li et al., 2015; Ververis et al., 2020), and lack of S1R exacerbates pathology of several neurological disorders (J. Hong, Wang, Zhang, Zhang, & Chen, 2017; Mavlyutov et al., 2013; Wang, Saul, Cui, Roon, & Smith, 2017). S1R is considered to be a potential drug target for treatment of neurodegenerative disorders and cancer (Herrando-Grabulosa, Gaja-Capdevila, Vela, & Navarro, 2020; Kim & Maher, 2017; D. A. Ryskamp et al., 2019). S1R binds multiple classes of drugs with nano- and sub-micromolar affinities (Maurice & Su, 2009). Based on their biological activity, S1R ligands are classified into agonists and antagonists (Maurice & Su, 2009). Signaling functions of S1R in cells are under intense investigation. The most prominent hypothesis is that under resting conditions S1R forms an inert complex with GRP78/BiP protein in the ER (Hayashi & Su, 2007). When activated by agonists or in conditions of cellular stress, S1R dissociates from BiP and is able to interact with client proteins (Hayashi & Su, 2007).

S1R preferentially localizes to MAMs (Hayashi & Su, 2007) and genetic knockout of S1R results in impaired MAM stability as evidenced by reduced number of contacts on EM and biochemical fractionation of MAM components (Watanabe et al., 2016). MAMs are cholesterol-enriched microdomains within ER membrane (Area-Gomez et al., 2012; Hayashi & Fujimoto, 2010; Montesinos et al., 2020), and previous studies demonstrated that S1R can directly interact with cholesterol and ceramides in MAMs (Hayashi & Fujimoto, 2010; Hayashi & Su, 2004; Palmer, Mahen, Schnell, Djamgoz, & Aydar, 2007). In the present study we tested the hypothesis that S1R association with cholesterol plays a critical role in organization of specialized lipid microdomains in the ER, including MAMs. Using two *in vitro* reconstitution systems, we demonstrate cholesterol-dependent clustering of S1R in lipid bilayers. We identified a novel cholesterol-binding site in the S1R sequence and demonstrated the importance of this site for S1R clustering. We further demonstrated that S1R agonists reduce S1R cluster size in the presence of cholesterol. Based on these results, we conclude that the main biological function of S1R in cells is related to its ability to organize and remodel cholesterol-enriched microdomains in the ER. Our conclusion is consistent with MAM defects observed in the S1R knockout mice (Watanabe et al., 2016).

## RESULTS

### S1R localizes to endoplasmic reticulum contact sites

To investigate S1R subcellular localization we transfected HEK293 cells with a S1R-GFP fusion protein construct and performed confocal imaging experiments. Previous reports indicate that intracellular localization of C-terminally tagged S1R is similar to that for endogenous receptor (Hayashi & Su, 2003b). To visualize the ER we co-transfected cells with mCherry-Sec61β (Zurek, Sparks, & Voeltz, 2011). In agreement with previous reports (Hayashi & Su, 2003a, 2003b), S1R-GFP formed puncta in the ER (Fig 1A, top panels). To determine if these puncta correspond to MAMs, we co-stained cells with a mitochondrial marker TOM20 and established that S1R-GFP puncta are frequently found in close opposition to mitochondria (Fig 1A, top panels). When an ALS-causing mutant S1RE102Q was expressed as a GFP-fusion, it was distributed uniformly in the ER and was not enriched in proximity to mitochondria (Fig 1A, lower panels). Cells with similar expression levels of S1R-GFP were compared in these experiments because high protein expression often led to S1R aggregation (data not shown). In order to increase the spatial resolution of our experiments we utilized an expansion microscopy (ExM) procedure that results in a 4.0-4.5 fold physical expansion of the specimen (Tillberg et al., 2016). HEK293 cells were transfected with S1R-GFP, stained with anti-GFP and anti-TOM20 antibodies and processed according to the pro-ExM procedure for imaging. On average, 12 ± 5 % of mCherry-Sec61β signal area overlapped with mitochondria, while higher degree of overlap was observed for S1R-GFP and mitochondria, with fractional area of 31 ± 7% (n (number of cells) = 5, p-value = 0.0034). These data confirmed that S1R-GFP is enriched in ER subdomains in close opposition to mitochondria (Fig 1B). These results suggested that S1R is enriched in MAM locations, consistent with previous reports (Hayashi & Su, 2007).

**Fig 1.**
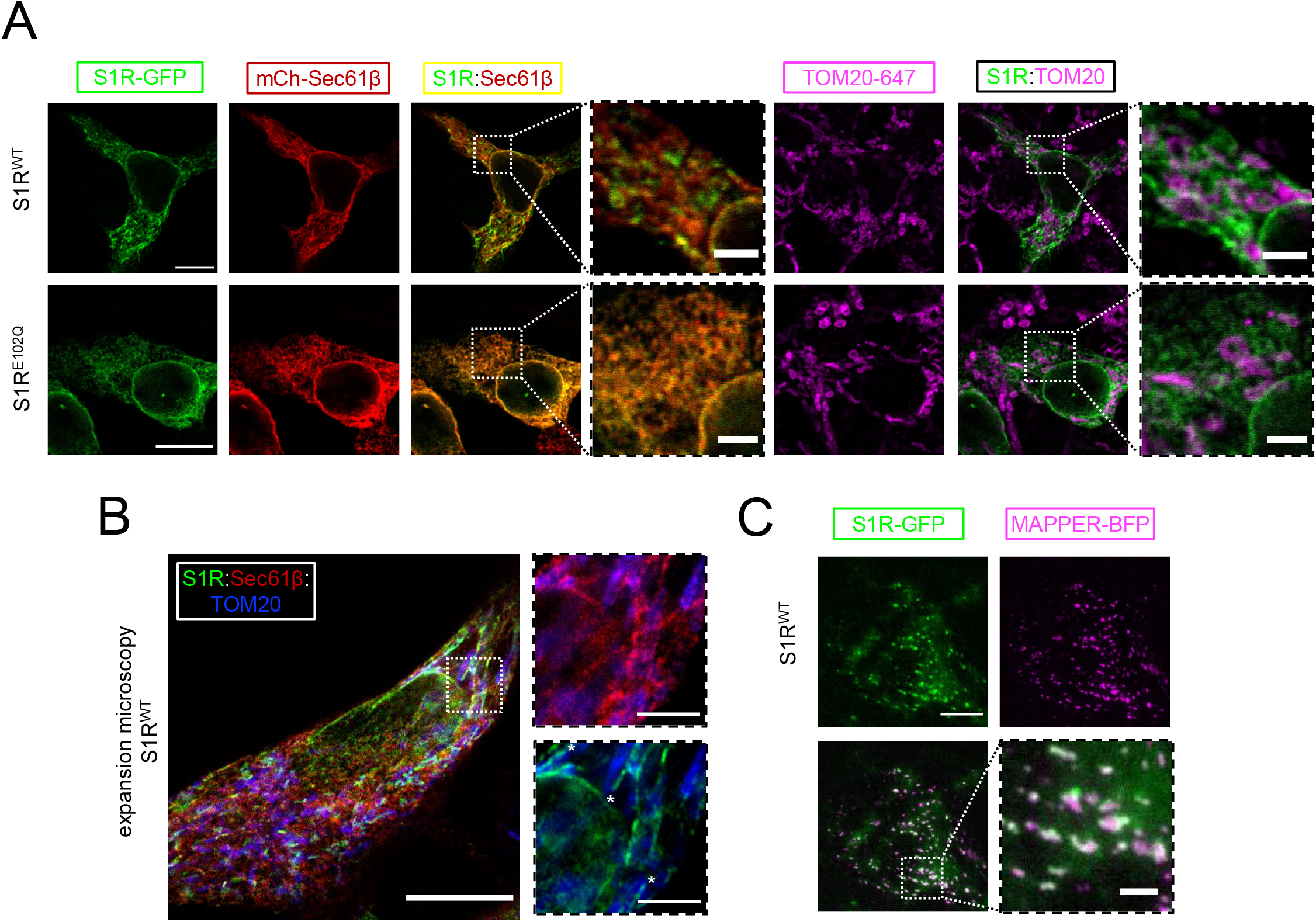
S1R targeting to ER contact sites in HEK293T cells. a. Intracellular distribution of wild type S1R-GFP (green, top panel) or ALS mutant variant E102Q-GFP (bottom panel) was visualized in HEK293 cells co-expressing ER marker mCherry-Sec61β (red) or immunostained with a mitochondrial marker TOM20 (magenta). Scale bar = 10 μm; = 2.5 μm for insets. b. Specimen was processed according to the protein retention expansion microscopy procedure, with S1R-GFP in green, mCherry-Sec61β in red and TOM20 staining. Insets show double staining of mCherry-Sec61β and TOM20 (top), S1R-GFP and TOM20 (bottom). Putative MAM compartments are labelled with asterisks. Scale bars = 5 μm, insets = 1.5 μm (real space). c. S1R localization at ER-PM junctions in HeLa cells visualized with a genetically-encoded maker MAPPER-BFP (magenta) and S1R-GFP (green). Cells were imaged using TIRF microscopy. Scale bars = 10 μm, = 2.5 μm inset.

We then asked if MAMs are the only inter-organelle contact sites that S1R localizes to. Previously S1R was shown to regulate store-operated Ca^2+^ entry (Srivats et al., 2016) and we reasoned that S1R may also be present at ER-plasma membrane (PM) junctions. To test this hypothesis, we co-expressed S1R-GFP with a genetically-encoded ER-PM marker MAPPER-BFP (Chang et al., 2013) in HeLa cells and imaged them using TIRF microscopy. As reported previously, MAPPER formed small punctate patterns corresponding to the ER membrane in close proximity to the PM visualized using TIRF microscopy (Fig 1C). S1R-GFP in the TIRF plane was also found in these same puncta (Fig 1C), in agreement with previous report (Srivats et al., 2016). The remaining diffuse fluorescence corresponds to S1R-GFP residing in the rest of the ER. Taken together, our results suggest that S1R localizes to ER contact sites such as MAMs and ER-PM junctions.

### S1R forms clusters in giant unilamellar vesicles *in vitro*

To investigate mechanisms responsible for S1R enrichment at ER subdomains, we performed a series of *in vitro* reconstitution experiments with giant unilamellar vesicles (GUV) (Fig 2A). NBD-labeled phosphatidylcholine (NBD-PC) was included in lipid mixtures used to generate GUVs for visualization. Full-length His-tagged S1R was expressed in Sf9 cells and purified to homogeneity by affinity chromatography, anion exchange chromatography and gel filtration chromagraphy (Fig 2B). After purification, His-S1R protein was covalently labelled with Alexa647 dye via NHS chemistry. To reconstitute S1R in the lipid membrane, large unilamellar vesicles (LUV) were prepared by extrusion, destabilized with detergent and mixed with purified S1R. Detergent was subsequently removed using BioBeads. S1R-containing LUVs were dehydrated on an agarose support and GUVs were formed by rehydration as previously described (Horger, Estes, Capone, & Mayer, 2009).

**Fig 2.**
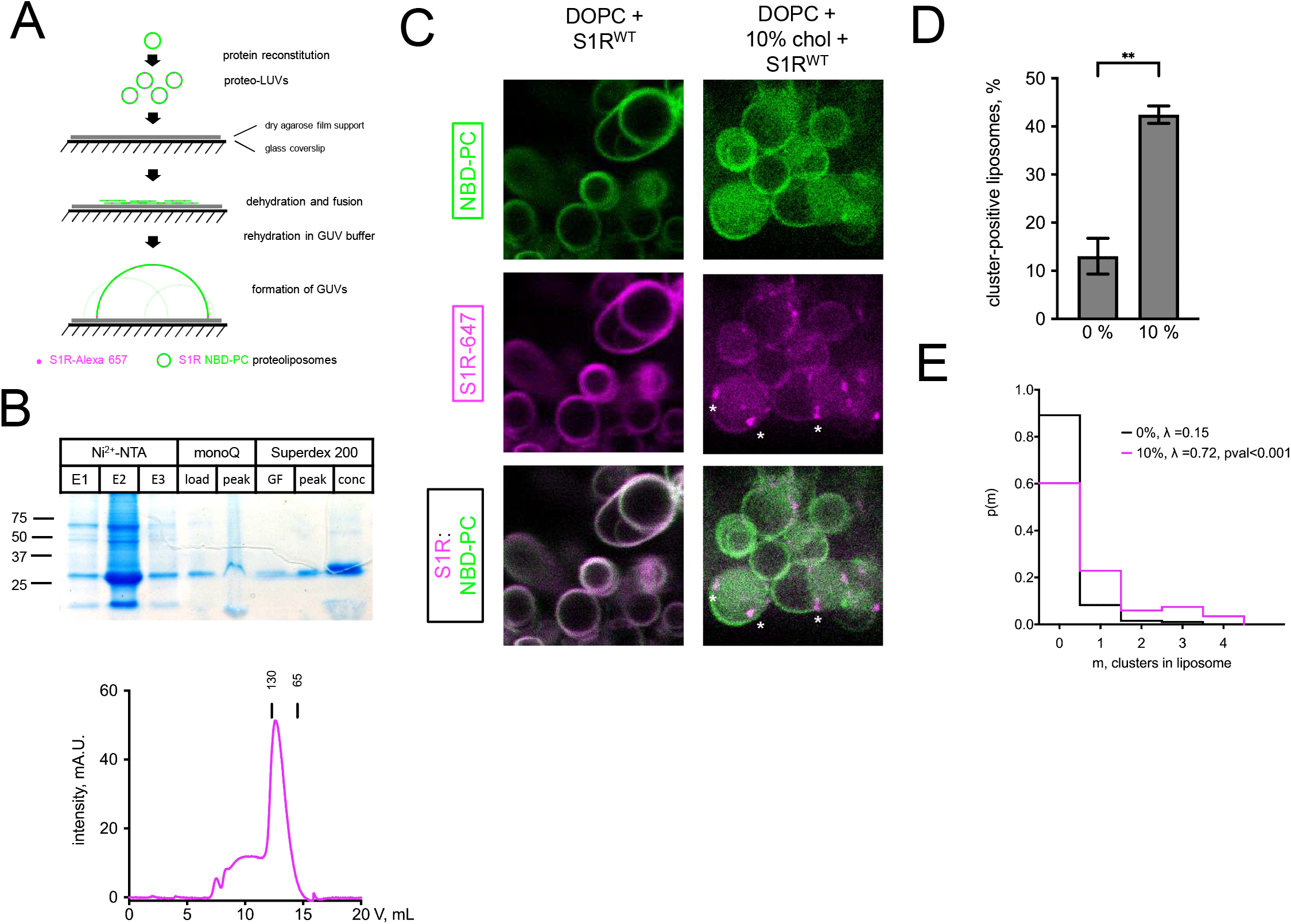
Cholesterol-dependent clustering of S1R in giant unilamellar vesicles (GUVs) a. GUV formation procedure. Purified S1R was labelled with Alexa647 and reconstituted into large unilamellar vesicles (LUV) to form proteoliposomes. After controlled dehydration of proteoliposomes on an agarose-covered coverslip, films were rehydrated in a salt buffer which resulted in the formation of micrometer-size GUVs. b. Biochemical characterization of purified S1R by SDS-PAGE analysis (top) and size-exclusion chromatography (bottom). c. Distribution of S1R-647 in cholesterol-free DOPC liposomes (left) and in the presence of 10% mol cholesterol (right), with membrane dye NBD-PC shown in green and S1R-647 in magenta. d. Average fraction of cluster-positive liposomes. In each experiment, 7 to 20 fields of view were analyzed. Data is mean ± SEM from n=3 independent experiments (0% condition: 411 liposomes, 10% condition: 528 liposomes). ** p=0.002 based on two-tailed t-test. e. Distribution of the number of clusters per liposome in the absence (black) and presence of 10 % mol cholesterol (magenta). Data was fitted with Poisson distribution to estimate the average number of clusters per liposome, λ (N1=173 liposomes for 0%, N2=201 for 10% mol cholesterol experimental condition). **** p-value <0.001 calculated based on Whitehead’s and C-test statistical tests.

Confocal imaging revealed that when S1R was reconstituted into liposomes composed of 18:1 PC (DOPC), it was distributed uniformly (Fig 2C, left panel) with occasional clustering observed at the sites of contacts between different GUVs (Sup Fig S1). In contrast, when vesicles also contained 10% cholesterol (molar ratio to DOPC), S1R was often clustered (Fig 2c, right panel, marked with asterisks). As cluster formation is a dynamic process, we made sure to prepare samples simultaneously and imaged them side by side. On average S1R clusters were observed in 13% ± 5% of DOPC liposomes (n = 3, 411 liposomes) and in 44% ± 6% of DOPC:cholesterol liposomes (n = 3, 528 liposomes) (Fig 2D). Frequently, the S1R coalesced into one large domain (more examples of S1R behavior in cholesterol-free and cholesterol-containing liposomes are presented in Suppl Fig S1). We used Poisson distribution to compare an average number of S1R clusters per liposome in each condition. We determined that the average number of clusters per liposome was significantly higher for GUVs containing 10% cholesterol (λ=0.15 for DOPC GUVs and λ=0.72 for cholesterol-containing liposomes, p<0.001, N1=173, N2=201) (Fig 2E). Similar results were obtained when 20% of cholesterol was used for formation of GUVs (data not shown). From these results we concluded that presence of cholesterol can promote clustering of S1R in the lipid membranes and propose that association with cholesterol plays a direct role in S1R-induced formation of ER membrane microdomains.

### Cholesterol binding motifs in S1R sequence

S1R was shown to interact with cholesterol, other sterols and sphingolipids in binding studies (Hayashi & Fujimoto, 2010; Hulce, Cognetta, Niphakis, Tully, & Cravatt, 2013; Palmer et al., 2007). Previous studies indicated two potential sites of S1R association with cholesterol - Y173 and Y201/Y206 (Palmer et al., 2007). However, in the S1R crystal structure (Schmidt et al., 2016) Y173 localizes adjacent to the S1R ligand binding domain and has no membrane contact, and Y201 and Y206 are located within the C-terminal membrane-adjacent amphipathic helices. Additional sequence analysis revealed tandem CARC-like (reverse sequence for cholesterol-recognition amino acid consensus) motifs R/K-X_1-5-_Y/F/W-X_1-5-_L/V (Fantini & Barrantes, 2013) in the transmembrane helix of S1R that spans amino acids 9-32 (Fig 3a). To test the importance of this motif, we generated S1R mutants by introducing a GGGG insertion within the CARC motif (S1R^4G^) or by mutating W9 and W11 to leucine residues (S1R^W9L/W11L^) (Fig 3A). We also generated Y173S and Y201S/Y206S mutants to test the potential importance of the previously reported cholesterol-binding motifs. In addition to the mutants describe above, we also generated a R7E/R8E mutant or deleted the double arginine motif altogether (ΔRR) (Fig 3A).

**Fig 3.**
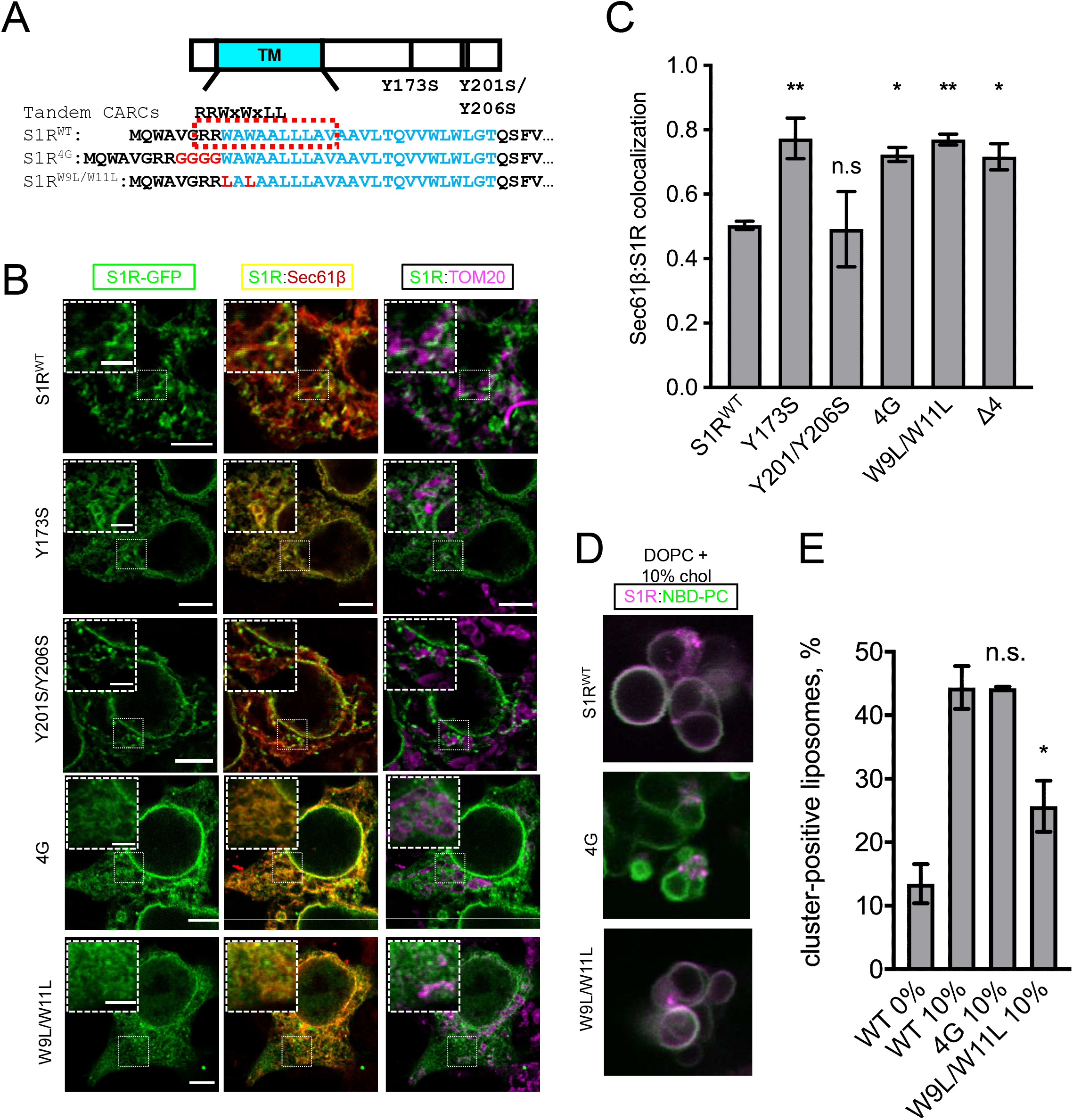
Cholesterol-binding motifs in the S1R sequence. a. Schematic representation of the S1R primary sequence with its transmembrane helix (TM) in cyan. Previously proposed cholesterol binding residues (Y173, Y201/Y206) are marked (Palmer et al., 2007). Sequence analysis identifies tandem CARC binding motifs in the TM region (marked in red). Mutations were introduced either by insertion of four-glycine repeat (4G) or by mutating critical tryptophan residues to leucine residues (W9L/W11L). b. Intracellular distribution of WT receptor and cholesterol-binding mutants in HEK293 cells, S1R-GFP in green, mCherry-Sec61β in red and anti-TOM20 in magenta. Scale bars = 5 μm, insets = 2.5 μm. c. Quantification of Mander’s colocalization coefficient between the mCherry-Sec61β and S1R-GFP for WT receptor and mutant forms. Data is mean ± SEM from n=3 independent experiments (n=2 for Y173S, Y201S/Y206S). p-values (n.s. p>0.05, * p<0.05, ** p<0.01): Y173S vs WT: p=0.008, Y201S/Y206S vs WT: p=0.999, 4G vs WT: p=0.014, W9L/W11L vs WT: p=0.004, Δ4 vs WT: p=0.017, based on ANOVA test with Dunnett’s post hoc test. d. Distribution of purified S1R-4G and S1R-W9L/W11L in cholesterol-containing GUVs with S1R-647 in magenta and NBD-PC in green. e. Average fraction of cluster-positive liposomes is shown for S1R, S1R-4G and S1R-W9L/W11L. Cluster-positive liposomes were quantified as in Fig 2D. Data (mean ± SEM) are from n=3 independent experiments. p-values (n.s. non-significant, * p-value < 0.05): WT 10% vs 4G 10%: p>0.99, W9LW11L 10% vs WT 10%: p=0.02 based on one-way ANOVA with Tukey’s post hoc test.

We first evaluated the effects of these mutations on S1R topology. We expressed wild type and mutant S1R fused to C-terminal GFP in HEK293 cells and stained them with anti-GFP antibodies following permeabilization of the plasma membrane with digitonin. When wild type S1R-GFP was expressed, we did not observe anti-GFP antibody staining (Suppl Fig S2), indicating that the C-terminus of S1R-GFP is located in the ER lumen, consistent with previous findings (Mavlyutov et al., 2017). Similar results were obtained with Y173S, Y201S/Y206S, 4G and W9L/W11L mutants (Suppl Fig S2). In contrast, intense anti-GFP antibody staining with observed in experiments with the R7E/R8E and ΔRR mutants (Suppl Fig S2), indicating that these proteins had reversed topology in the ER membrane. Therefore, we excluded these mutants from further analysis.

Wild type and mutant S1R were expressed in HEK293 cells and their distribution was analyzed by confocal microscopy. Wild type S1R and the S1R-Y201S/Y206S mutant each formed puncta in the ER (Fig 3B). In contrast, the S1R distribution in the ER was diffuse for the S1R-Y173S, S1R-4G and S1R-W9L/W11L mutants (Fig 3B). By performing TOM20-staining experiments we confirmed that the puncta observed with the wild type S1R and Y201S/Y206S mutant correspond to MAMs (Fig 3B). In order to obtain a quantitative measure of the S1R distribution in the ER network, we calculated the Mander’s overlap coefficient between mCherry-Sec61β and S1R-GFP. Mander’s coefficient measures fractional overlap between two channels and varies from 0 (no correlation) to 1 (complete correlation). For wild type S1R, the Mander’s coefficient was M=0.50±0.02 (n=3, 25 cells), indicating that not all mCherry-Sec61β signal co-localized with the GFP signal in S1R clusters (Fig 3C). In contrast, the Mander’s coefficient was equal to 0.77±0.09 (n =2, 11 cells) for S1R-Y173S, 0.72±0.04 (n=3, 30 cells) for S1R-4G, and 0.77±0.03 (n=3, 17 cells) for S1R-W9L/W11L (Fig 3C), reflecting more diffuse distribution patterns. The Mander’s coefficient for the S1R-Y201S/Y206S mutant was equal to 0.49±0.16 (n=2, 12 cells) (Fig 3C), similar to the wild type. Based on these results we conclude that the newly identified CARC motif is important for S1R targeting to MAMs and that the Y201/Y206 motif is dispensable. Effects of Y173 mutation may be related to misfolding of S1R as this residue has no membrane contact in the S1R crystal structure (Schmidt et al., 2016).

To further validate the importance of the CARC motif in S1R cholesterol-dependent clustering, we expressed and purified S1R-4G and S1R-W9L/W11L mutant proteins and performed GUV reconstitution experiments in the presence of 10% cholesterol. The S1R-W9L/W11L mutant did not form clusters (Fig 3D, 3E), consistent with the diffuse distribution of this mutant in cells (Fig 3B, 3C). However, S1R-4G still formed clusters in when reconstituted in GUVs, similar to wild type S1R (Fig 3D, 3E). This was in contrast to the diffuse distribution of the S1R-4G mutant in cells (Fig 3B, 3C). To explain these results we reasoned that the W9L/W11L mutation abolished S1R association with cholesterol, but the 4G mutation only weakened it as this mutation still contains the CARC consensus sequence (Fantini & Barrantes, 2013). ER cholesterol levels in cells are less than 10% (Radhakrishnan et al., 2008), insufficient for binding to the S1R-4G but sufficient for binding to the wild type S1R.

### S1R localizes to, and can generate, thick lipid domains

Our results suggest that S1R is clustered in cholesterol-rich microdomains in the ER (Fig 3). It is established that cholesterol-rich phospholipid mixtures have more ordered acyl chains and larger hydrophobic thickness (de Meyer & Smit, 2009). Interestingly, according to the crystal structure and hydrophobicity analysis the length of S1R transmembrane domain is 24 a.a (Fig 3A) (Schmidt et al., 2016), longer than the typical 20 a.a TM length of ER-resident proteins (Sharpe, Stevens, & Munro, 2010). The length of the S1R TM domain is estimated to be 36.9 Å and should have a positive hydrophobic mismatch when compared to a DOPC bilayer, which has a hydrophobic thickness of 26.8 Å (Kucerka, Tristram-Nagle, & Nagle, 2005). In contrast, a cholesterol-rich DOPC bilayer has hydrophobic thickness of 36.0 Å, matching well with the S1R TM length (Milovanovic & Jahn, 2015). It is therefore possible that S1R clustering in ER membrane is influenced by local changes in membrane thickness, as has been described for targeting of plasma membrane proteins to lipid rafts (Lorent et al., 2017). To test whether the length of the transmembrane domain of S1R also contributes to clustering, we deleted four amino acid residues in its transmembrane region (Fig 4A). When S1R^Δ4^-GFP protein was expressed in HEK293T cells, it displayed a more diffuse distribution in the ER when compared to wild type S1R (Fig 4A). The Mander’s coefficient for this mutant is 0.72±0.07 (n=3, 27 cells), compared to M=0.50±0.02 for wild-type protein (Fig 3C). S1R^Δ4^-GFP also did not show MAM localization based on TOM20 staining (Fig 4A).

**Fig 4.**
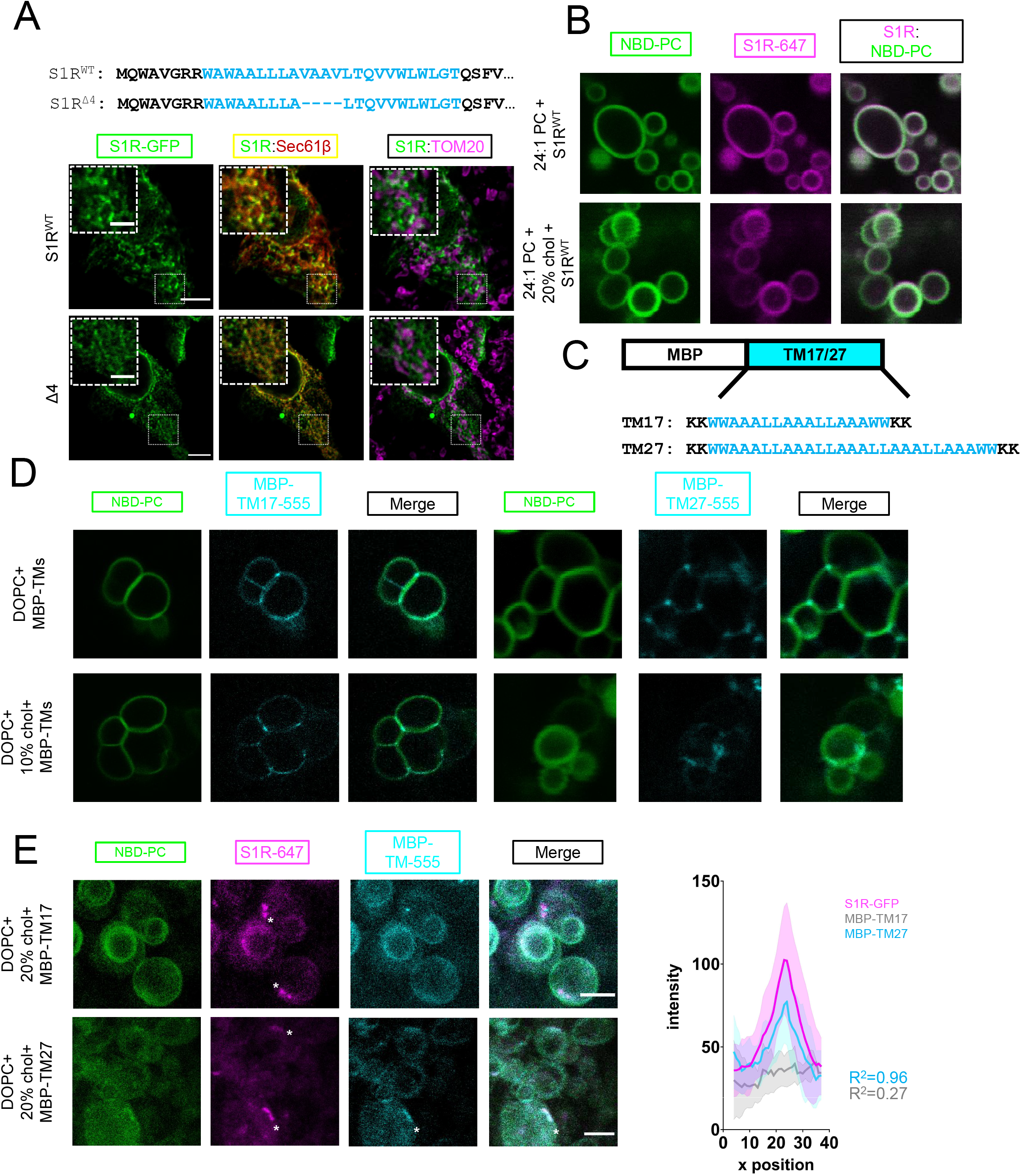
Importance of membrane bilayer thickness for S1R cluster formation. a. Deletion of the four amino acid stretch from the S1R TM domain (top) and intracellular localization of full-length (WT) and short mutant (Δ4) in HEK293 cells. S1R-GFP in green, mCherry-Sec61β in red and anti-TOM20 in magenta. Scale bars = 10 μm, insets = 2.5 μm. Mander’s colocalization coefficient for the Δ4 mutant in plotted on Fig 3C. b. Distribution of S1R-647 (magenta) in 24:1 PC GUVs (NBD-PC in green) in the absence (top) or presence of 20 % mol cholesterol (bottom). c. Construct design and primary amino acid sequences of MBP-TM17 and -TM27 proteins. Synthetic transmembrane domain in shown in cyan. Construct design based on (Kaiser et al., 2011). d. Lateral distribution of MBP-TM17 (cyan, left panel) and MBP-TM27 (cyan, right panel) in DOPC GUVs (NBD-PC, green) in the absence (top) of presence (bottom) of 10 % mol cholesterol. e. Distribution of S1R-647 (magenta) co-reconstituted together with MBP-TM17-555 (cyan, top panel) or MBP-TM27-555 (bottom panel) in DOPC GUVs in the presence of cholesterol (NBD-PC in green). Inset shows fluorescent intensity distribution across a S1R cluster. Pearson correlation coefficients were calculated for S1R-647 and MBP-TM17-555 (R2=0.27) or -TM27-555 (R2=0.96) for n=5 clusters.

To test a role of positive hydrophobic mismatch in S1R clustering, we reconstituted purified Alexa647-labeled S1R in GUV membranes composed of 24:1 PC (DNPC) lipids that have a hydrophobic thickness of 35.5 Å. S1R did not form clusters in DNPC GUVs either in the absence or in the presence of 20% cholesterol (Fig 4B). From these results we concluded that local increase in membrane thickness contributes to S1R targeting to cholesterol-enriched microdomains in the ER. To measure local membrane thickness in these microdomains, we utilized two designed molecular rulers that report on transmembrane thickness - MBP-TM17 with a short transmembrane helix and MBP-TM27 with a longer transmembrane helix (Fig 4C). The amino acid sequences of TM17 and TM27 were designed based on synthetic peptides previously used in GUV studies of membrane thickness (Kaiser et al., 2011). Similar WALP-GFP fusion proteins were successfully used for measurements of ER lipid heterogeneity in yeast cells (Prasad, Sliwa-Gonzalez, & Barral, 2020). The hydrophobic length of MBP-TM17, calculated using 1.5 Å rise per residue (Hildebrand, Preissner, & Frommel, 2004), matches well with the hydrophobic thickness of a DOPC bilayer (25.5 Å, and 26.8 Å, respectively) (Kucerka et al., 2005). The calculated hydrophobic length of TM27 is longer (40.5 Å) and matches well to a lipid bilayer with longer acyl chains or a bilayer with high cholesterol content (DOPC + 30 - 40 mol % cholesterol, 36.0 Å) (Milovanovic et al., 2015). MBP-TM17 and MBP-TM27 proteins were expressed in *E. coli*, purified using affinity and size exclusion chromatography, and then covalently labelled with Alexa555 dye (Suppl Fig S3). The MBP-TM17 and MBP-TM27 were reconstituted into DOPC GUVs in the absence or in the presence of 10% cholesterol. We found that in the cholesterol-free vesicles MBP-TM17 was distributed largely uniformly in the GUV membrane while MBP-TM27 was enriched at the junctions between individual GUVs (Fig 4D, top panel). Cholesterol had little effect on the distribution of MBP-TMs (Fig 4D, bottom panel). We hypothesized that MBP-TM27 localized to the GUV junction sites based on a hydrophobic matching mechanism. Interestingly, clustering of S1R at the sites of contacts between different GUVs was occasionally observed in the absence of cholesterol (Fig 2C, left panel), most likely due to increase in local membrane thickness at these sites. When these rulers were reconstituted together with S1R in the presence of 20% cholesterol, they displayed a different behavior. MBP-TM17 remained largely diffuse (Fig 4E, top panel) but MBP-TM27 formed clusters that co-localized with S1R clusters (Fig 4E, bottom panel). Analysis of intensity distribution of S1R-647 and MBP-TM signal showed high degree of correlation for S1R-647 and MBP-TM27 with calculated Pearson’s correlation coefficient of R2=0.96 (Fig 4 E, inset). Importantly, this behavior was observed for an artificially designed MBP-TM27 protein, suggesting that recruitment and clustering events were driven mainly by lipid-protein interactions. These results suggested that in the presence of cholesterol S1R induces formation of membrane microdomains with increased local thickness.

### The dynamic organization of S1R clusters in double lipid supported bilayers

Based on their biological activity, agonists and antagonists of S1R have been described (Maurice & Su, 2009). Previous research indicated that S1Rs can redistribute in the ER upon ligand stimulation (Hayashi & Su, 2003a, 2007). In the next series of experiments, we sought to understand whether S1R ligands can analogously effect the formation and stability of S1R clusters *in vitro*. It is technically difficult to measure time-resolved dynamics of protein clustering in the membrane of GUVs using standard confocal microscopy. Thus, we used total internal reflection fluorescence (TIRF) microscopy and supported lipid bilayers (SLBs), a system commonly employed to measure the dynamics of membrane-associated proteins (X. Su et al., 2016). SLBs are typically formed on a glass or mica surface. But in this format transmembrane proteins bind to the underlying glass/mica and become immobilized. Recently, several groups reported formation of polyethylene-glycol (PEG)-cushioned bilayers that retained mobility of transmembrane proteins (Pace et al., 2015; Richards et al., 2016; Wong et al., 2019). When we attempted to form supported or cushioned bilayers with reconstituted S1R, we observed that protein and lipids were static, presumably due to interaction with the glass surface (data not shown). To overcome these limitations, we modified the bilayer formation procedure to generate double supported lipid bilayers (DSLBs). In DSLBs, PEGylated lipids are used as supports between two separate bilayers: a lower bilayer that contacts the glass surface is composed of DOPC and 1 mol % DSPE-PEG2000-biotin lipids and an upper bilayer that rests on the first and is composed of lipids and purified transmembrane proteins. To build the DSLB, the first bilayer is formed using small unilamellar vesicles (SUVs) prepared by freeze-thaw of DOPC and 1 mol % DSPE-PEG2000-biotin lipids by fusion and spreading on a clean glass surface (Fig 5A). The second bilayer is formed by deposition of extruded LUVs containing 0.1% NBD-PC as a membrane dye that is used to measure membrane lipid fluidity by TIRF microscopy (Fig 5A). Fluorescence recovery after photobleaching (FRAP) of NBD-PC indicated that lipids in the upper bilayer are highly mobile (Suppl Figs S4A, S4B). To incorporate protein in the DSLB, purified S1R-Alexa647 was first reconstituted in LUVs using the same approach as for the GUV studies. Protein-loaded LUVs were then applied to the pre-formed lower DOPC bilayer and allowed to spread and form a continuous upper bilayer (Fig 5A). The second bilayer was within the TIRF penetration depth (100 nm). FRAP experiments indicated that S1R molecules in this system are mobile and can quickly diffuse laterally in the membrane (Suppl Fig S3C).

**Fig 5.**
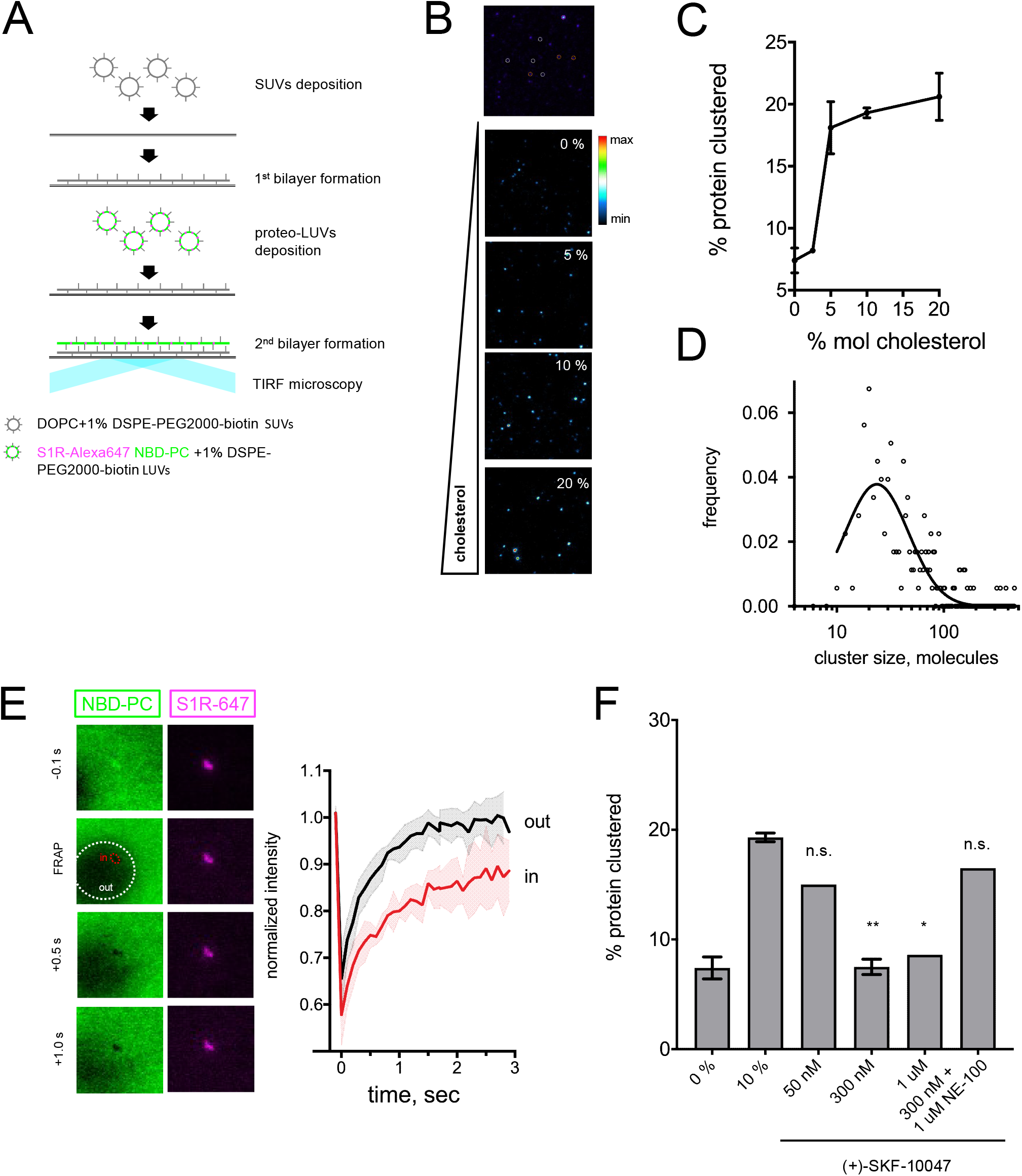
Visualization of S1R clusters in double supported lipid bilayers. a. The double supported lipid bilayer (DSLB) technique. Small unilamellar vesicles (SUV) vesicles prepared from DOPC and 1 % pegylated lipids are deposited on a glass surface. S1R-Alexa647 (magenta) is reconstituted into LUVs doped with 1% pegylated lipids and NBD-PC dye (green). Proteoliposomes are deposited on a pre-formed first bilayer. Bilayer is imaged using TIRF microscopy. b. Detection of S1R clusters in DSLB (top image). Trimeric receptor (orange circles), bigger assemblies (white circles) and large clusters (magenta) can be identified in DSLBs. Cholesterol-dependent clustering of S1R-647 in DSLB. Heat map coloring scheme was used for better visibility. c. Quantification of the clustered fraction versus membrane cholesterol concentration. In each well, at least ten fields of view were quantified, and experiments were repeated independently n=3 times (n=1 for 2.5%, n=2 for 5.0%). d. Size distribution of higher-order S1R clusters (more than ten molecules) in DSLB at 20% mol cholesterol based on their fluorescent intensity (open circles) and lognormal fit of the data (solid line). e. Lipid dynamics in S1R clusters measured by FRAP. A small area of the bilayer was bleached, and fluorescence recovery of NBD-PC dye was monitored outside (out, white area) and inside (in, red area) of a S1R cluster. Normalized FRAP curves (red within the cluster, black for outside bilayer) are plotted from n=3 FRAP experiments. f. Ligand effects of S1R agonist and antagonist on S1R clustering in DSLB. Increasing concentrations (50 nM to 1 μM) of a selective S1R agonist (+)-SKF-10047 were added to the wells during the second bilayer formation step (2 h) and clustered fraction was quantified. S1R antagonist NE-100 at 1 μM concentration was added together with 300 nM (+)-SKF-100047. Data are mean ± SEM. In each experiment, at least ten fields of view were analyzed. p-values (n.s. p>0.05, * p-value<0.05, ** p<0.01): 0% vs 10%: p=0.007, 50 nM vs 10%: p=0.187, 300 nM vs 10%: p=0.007, 1 μM vs 10%: p=0.173, 300 nM + 1 μM NE-100: p=0.43 based on one-way ANOVA with Tukey’s post hoc test.

We discovered that the S1R distribution in DSLBs was heterogeneous and based on cluster intensity, small and large clusters can be identified (Fig 5B). Small clusters of S1R were highly mobile in the DSLB membrane, whereas large clusters were relatively static. To investigate effects of cholesterol on S1R clustering, we incorporated S1R-Alexa647 into DSLBs containing increasing amounts of cholesterol (Fig 5B). In agreement with GUV data (Fig 2), we found that increased cholesterol in DSLBs caused S1R to cooperatively assemble into large clusters (Fig 5B). The threshold for S1R clustering was in a narrow range between 2.5-5.0 mol % (Fig 5C), consistent with a typical ER cholesterol content (Radhakrishnan et al., 2008). To determine the number of S1R molecules in each of these clusters, we divided the integrated intensity of each cluster by the intensity of a mono-labelled His-pLAT-647 protein molecule (X. Su et al., 2016). This analysis revealed that small S1R clusters most likely correspond to S1R trimers, containing an estimated average of 2.65 ± 0.58 (n = 47) molecules, consistent with the crystal structure of the protein (Schmidt et al., 2016). The number of S1R molecules in the large clusters was widely distributed in a range between tens and hundreds of molecules, with a mean value of 38 molecules per cluster (Fig 5D). To measure the relative lateral mobility of the lipids, we performed FRAP experiments on NBD-PC within S1R clusters and in the surrounding the membrane (in the presence of 20% cholesterol). Following photobleaching, the recovery of NBD-PC signal was significantly slower within S1R clusters (Fig 5E). These data suggest that lateral mobility of the lipids within S1R clusters is reduced, consistent with known properties of lipid rafts in the plasma membrane (Sezgin, Levental, Mayor, & Eggeling, 2017).

We next evaluated the effects of S1R ligands on S1R clustering. These experiments were performed in the presence of 10% cholesterol in the DSLB. Addition of selective S1R agonist (+)-SKF-10047 at the time of the second bilayer formation reduced the number of large S1R clusters (Fig 5F). The effects of (+)-SKF-10047 on S1R clustering were concentration-dependent, with half-maximal effect observed at 50 nM (Fig 5F). The effects of (+)-SKF-10047 were blocked by addition of S1R antagonist NE-100 (Fig 5F). These results suggested that S1R agonists prevent formation of S1R clusters or promote disassembly of S1R clusters in the membrane and that these effects can be blocked by S1R antagonists. Our results are consistent with previous biochemical studies that demonstrated that the proportion of S1R multimers formed in cells was decreased by the agonists (+)-pentazocine and PRE-084 but increased by the antagonists CM304, haloperidol and NE-100 (W. C. Hong, 2020; W. C. Hong et al., 2017). These results are also consistent with effects of (+)-pentazocine and haloperidol described in FRET experiments performed with cells transfected with fluorescently tagged S1Rs (Mishra et al., 2015). In contrast, it has been reported that S1R agonists such as (+)-pentazocine and PRE-084 stabilized the oligomeric state of purified S1R in the presence of detergents (Gromek et al., 2014), suggesting that biochemical properties of S1R in the membrane and in detergent may differ from each other. Another potential reason for discrepancy is that previous study (Gromek et al., 2014) was focused on the analysis of smaller tetrameric species, while our analysis measured larger assemblies of at least ten S1R molecules.

### S1R sustains UPR signaling by recruiting IRE1α

To examine biological relevance of S1R-formed lipid microdomains, we focused on analysis of signaling by inositol-requiring enzyme 1α (IRE1α). S1R and IRE1α localize in a close proximity to each other in the ER, but do not necessarily interact directly (Rosen et al., 2019). In addition to protein-protein interactions, IRE1α oligomerization and stress-response activity can be also modulated by the surrounding lipid environment (Cho et al., 2019; Halbleib et al., 2017). To confirm colocalization of S1R and IRE1α in cells, we utilized a proximity labelling approach (Hung et al., 2016). HEK293 cells were transiently transfected with S1R-APEX2 protein or control ER-targeted constructs APEX2-KDEL or Sec61β-APEX2. Following transfection, cells were incubated with biotin-phenol and exposed to H_2_O_2_ for a short period of time to induce biotinylation of proteins in proximity to APEX2. Cell lysates were collected, and biotinylated proteins were pulled down using streptavidin-agarose. The eluate containing streptavidin-agarose beads bound by biotinylated proteins was analyzed by Western blot. Significantly more biotinylated IRE1α was pulled down from the cells expressed S1R-APEX2 than from the cells expressing APEX2-KDEL or Sec61β-APEX2 control constructs (Fig 6A). These results suggest close proximity of IRE1α and S1R-APEX2, consistent with previous studies (Mori, Hayashi, Hayashi, & Su, 2013; Rosen et al., 2019). Short-term (2 h) stress induction with 1 μM thapsigargin (Tg) had no measurable effect on co-localization of S1R and IRE1α.

**Fig 6.**
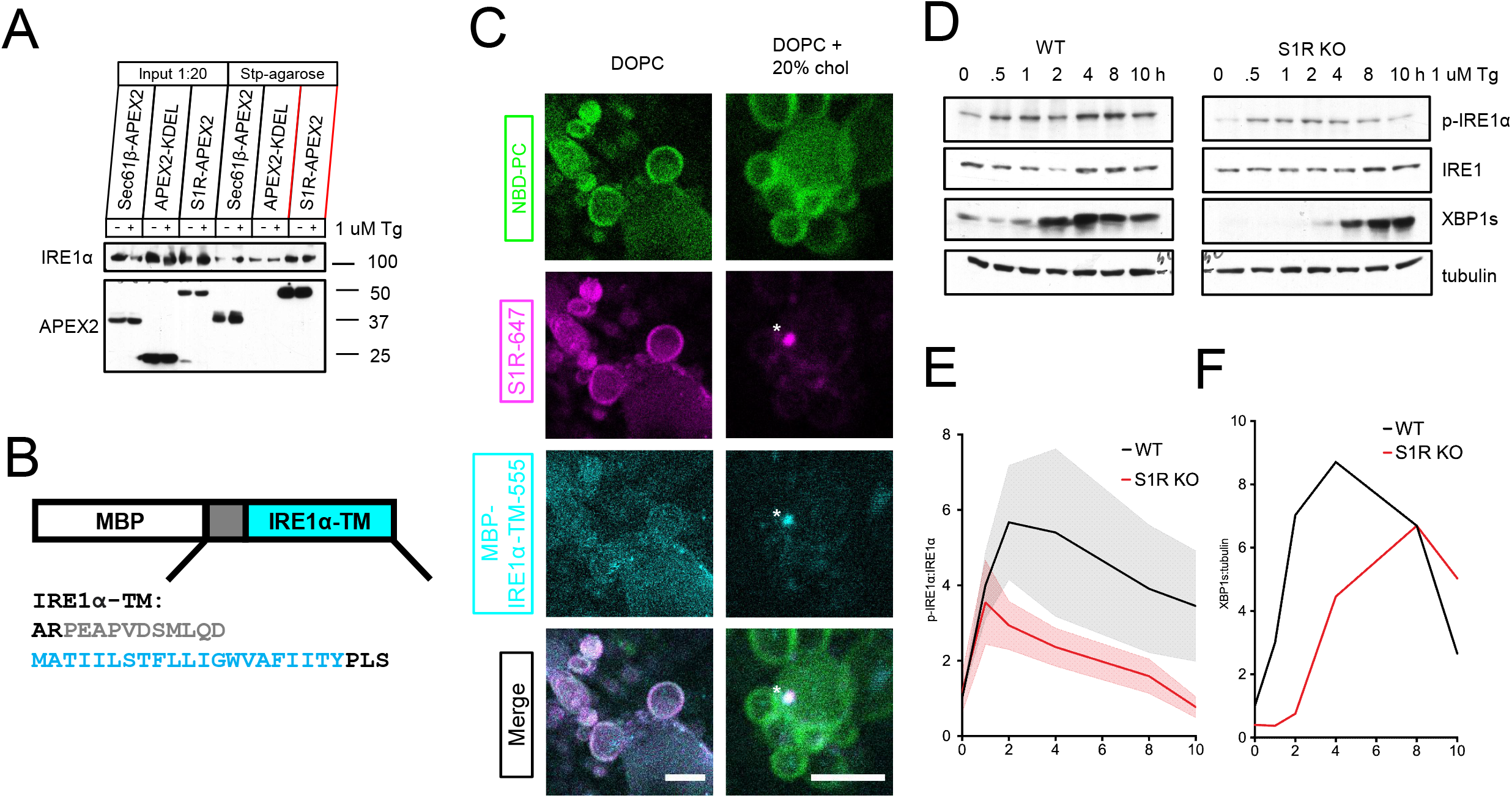
Importance of S1R for IRE1α-mediated signaling. a. Analysis of proximity labelling of IRE1α. HEK293 cells expressed S1R-APEX2 and Sec61β-APEX2 or APEX2-KDEL as controls. Cells were incubated with biotin-phenol and proteins were labeled by a 1-minute incubation with H_2_O_2_. Biotinylated proteins were pulled down and analyzed by Western blot. b. Construct design and primary amino acid sequence of a minimal sensor derived from human IRE1α. TM helix of IRE1α is shown in cyan and the adjacent amphipathic helix is in grey. Construct design based on (Cho et al., 2019). c. Distribution of S1R-647 (magenta) and co-reconstituted MBP-IRE1α-TM labelled with Alexa555 (cyan) in GUV (NBD-PC in green) in the absence (left) and presence of 20% mol cholesterol. Scale bars = 10 μm (left), = 5 μm (right). d. Time-dependence of the thapsigargin-induced IRE1α response in WT and S1R KO HEK293 cells. Western blot is representative of n=3 independent experiments. e. and f. Quantification results of the Western blot shown above. p-IRE1α levels were normalized to the total level of IRE1α and plotted as mean (solid line) ± SEM (shaded) for WT (black) and S1R KO cells (red), XBP1s levels were normalized to tubulin.

To test whether S1R co-assembles with IRE1α in GUV membranes, we used a similar approach that we utilized for the MBP-TM rulers (Fig 4). For these experiments we generated an expression construct MBP-IRE1α-TM that consists of the MBP protein fused to the transmembrane helix of human IRE1α and the upstream amphipathic helix (433-464 a.a) (Fig 6B) (Cho et al., 2019). MBP-IRE1α-TM protein was expressed in bacteria, purified and labelled with Alexa555 (Suppl Fig S5). MBP-IRE1α-TM-555 protein was co-reconstituted together with S1R-647 in GUVs and imaged by fluorescence confocal microscopy. In the absence of cholesterol, the distributions of MBP-IRE1α-TM and S1R were uniform (Fig 6C, left panel). However, in the presence of 20% cholesterol, MBP-IRE1α-TM was recruited to S1R-positive clusters (Fig 6C, right panel). These results suggest that the transmembrane domain of IRE1α partitions into a lipid microenvironment established by S1R in the presence of cholesterol.

To examine the functional relevance of IRE1α localization to S1R microdomains in the ER, we compared IRE1α-mediated signaling in wild type and S1R knockout (KO) HEK293 cells (D. Ryskamp et al., 2017). We induced the unfolded protein response (UPR) by addition of 1 μM Tg and monitored levels of IRE1α phosphorylation by Western blot using a phospho-IRE1α antibody. We determined that under resting conditions the level of phosphorylated pIRE1α was lower in S1R KO cells when compared to wild type cells (Fig 6D, first lanes). Under ER stress conditions the same peak level of IRE1α phosphorylation could not be achieved in S1R KO cells (Fig 6D, 6E), suggesting impaired IRE1α signaling. To further examine the activity of IRE1α, we quantified the levels of the XBP1s protein, a known downstream effector of IRE1α (Calfon et al., 2002). We found that production of XBP1s was significantly delayed in S1R KO cells when compared to wild type cells (Fig 6D, 6F). These results suggest that recruitment of IRE1α to S1R-organized microdomains in the ER facilitates IRE1α-mediated signaling. Our conclusions are consistent with previously reported inhibition of IRE1α activity in cells transfected with S1R RNAi (Mori et al., 2013).

## DISCUSSION

S1R modulates many physiological processes, such as cell excitability, transcriptional activity, Ca^2+^ homeostasis, stress response and autophagy (Christ, Huesmann, Nagel, Kern, & Behl, 2019; Couly et al., 2020; Hayashi, 2019; Kourrich, 2017; Maurice & Goguadze, 2017; D. A. Ryskamp et al., 2019). However, S1R-mediated signal transduction differs from canonical second messenger-coupled transmembrane receptor signaling and S1R is often referred to as a “ligand-gated molecular chaperone” (Hayashi, 2019; Nguyen et al., 2017; T. P. Su et al., 2010). In the seminal paper by Hayashi and Su (Hayashi & Su, 2007), S1R was proposed to have chaperone-like properties at MAMs. However, the molecular mechanism of chaperone activity of S1R remains unexplained because S1R lacks structural similarity with known chaperones or extensive protein interaction interfaces. Based on our results, we propose that the biological activity S1R in cells can be explained by the ability of this protein to form cholesterol-enriched microdomains in the ER, therefore acting as ER lipid scaffolding protein. A role of S1R in organization and remodeling of lipid raft microdomains in the plasma membrane has been proposed previously based on cell biological studies (Palmer et al., 2007; Vollrath et al., 2014). Our data further suggest that such microdomains have increased local membrane thickness, providing a favorable environment for recruitment of ER proteins with longer transmembrane domains (Lorent et al., 2017). Increased local cholesterol concentration and membrane thickness can modulate activity of ER proteins recruited to these microdomains. Our hypothesis may explain how a small protein such as S1R is able to modulate activity of almost a hundred effector proteins (Couly et al., 2020; Delprat, Crouzier, Su, & Maurice, 2020; Kourrich, Su, Fujimoto, & Bonci, 2012; D. A. Ryskamp et al., 2019; Schmidt & Kruse, 2019). We reason that activity of these proteins could be affected by changes in local lipid microenvironment and not only via protein-protein interactions with the S1R. By performing experiments in reduced reconstitution systems, we have been able to demonstrate direct effects of cholesterol on S1R clustering (Figs 2 and 5). Previous studies indicated two potential sites of S1R association with cholesterol - Y173 and Y201/Y206 (Palmer et al., 2007). Our studies suggest that Y201 and Y206 residues are dispensable for cholesterol-mediated S1R clustering (Fig 3). Moreover, we identified a tandem CARC-like motif (Fantini & Barrantes, 2013) within the transmembrane region of S1R (Fig 3A). Mutations of this motif affected S1R clustering in cells and *in vitro*, suggesting its importance for S1R interaction with cholesterol. The threshold for formation of S1R clusters was in a narrow range between 2.5-5.0 mol % (Fig 5C). Similar cholesterol dependence was previously described for SREBP-2 activation, with half-max response at 4.5% mol of ER cholesterol (Radhakrishnan et al., 2008), indicating that S1R affinity for cholesterol is within physiological range of ER cholesterol concentrations.

A recent study demonstrated that micrometer-sized large intracellular vesicles exhibit phase-separation behavior at contact sites between ER and mitochondria, plasma membrane and lipid droplets membranes (King et al., 2020), similar to phase separation observed for giant plasma membrane-derived vesicles (Levental, Grzybek, & Simons, 2011). It has been shown that MAMs are enriched in cholesterol and ceramides (Area-Gomez et al., 2012; Hayashi & Fujimoto, 2010; Hayashi & Su, 2003a). Our findings suggest that S1R can contribute to stabilization and/or formation of these cholesterol-rich lipid microdomains in the ER membrane. This conclusion is consistent with MAM defects observed in S1R knockout mice (Watanabe et al., 2016), with earlier analysis of S1R targeting to detergent-resistant domains in the ER (Hayashi & Fujimoto, 2010), and with previous suggestions that S1R contribute to stability of lipid rafts in the plasma membrane (Palmer et al., 2007; Vollrath et al., 2014). The lipid regulation of PM proteins is a well-known phenomenon (Rosenhouse-Dantsker, Mehta, & Levitan, 2012). It was shown in direct and indirect experiments that activities of several ER channels including ryanodine receptors, sarco/endoplasmic reticulum Ca^2+^-ATPase and inositol-1,4,5-triphosphate receptors can be modulated by cholesterol content, lipid packing or membrane thickness (Cannon et al., 2003; Gustavsson, Traaseth, & Veglia, 2011; Y. Li et al., 2004; Madden, Chapman, & Quinn, 1979; Sano et al., 2009). Components of the gamma secretase complex reside at the MAM and transmembrane region of APP contains cholesterol-binding motifs (Montesinos et al., 2020; Pera et al., 2017), suggesting modulatory effects of cholesterol on APP processing. Several ER stress sensors, including IRE1α, can sense membrane saturation and be activated without protein unfolding in the ER lumen (Ballweg et al., 2020; Cho et al., 2019; Halbleib et al., 2017). In our experiments we demonstrated that a minimal sensor derived from IRE1α is recruited to S1R clusters (Fig 6C) and S1R knockout impaired IRE1α-mediated signaling (Fig 6D, 6E) in agreement with the previous report (Mori et al., 2013), providing a mechanistic explanation for S1R-mediated potentiation of the IRE1α response.

S1R can exist in oligomeric form (Gromek et al., 2014; Mishra et al., 2015). Originally, a photoaffinity probe labelled high-molecular weight oligomers in rat liver microsomes (Pal et al., 2007). High-molecular weight species up to 400 kDa were later detected in membrane preparations using various biochemical approaches (Yano et al., 2018; Yano, Liu, Naing, & Shi, 2019) and a similar wide molecular weight distribution was observed for the recombinant protein (Schmidt et al., 2016). The functional relevance of the oligomerization is however not clear. In our experiments we observed S1R in the form of continuous molecular weight distribution ranging from trimers to large (10-100 subunits) clusters (Fig 5). Formation of large clusters was facilitated by the presence of cholesterol. S1R agonists disrupted large S1R clusters in cholesterol-containing lipid bilayers in a concentration-dependent manner (Fig 5F). Our results are consistent with ligand effects observed by native electrophoresis from cell membranes and S1R-BiP association (Hayashi & Su, 2007; W. C.Hong, 2020; W. C. Hong et al., 2017). Also in agreement with our findings, S1R agonist SKF10047 resulted in destabilization of lipid rafts in the plasma membrane of cancer cells (Palmer et al., 2007)

Mutations in S1R lead to a juvenile form of ALS (Al-Saif et al., 2011) and distal hereditary motor neuropathy (Almendra et al., 2018; Gregianin et al., 2016; Horga et al., 2016; X. Li et al., 2015; Ververis et al., 2020). We demonstrated that ALS-causing E102Q variant acts as a loss-of-function mutant that disrupts S1R targeting to MAMs (Fig 1A). S1R is considered to be a potential drug target for treatment of neurodegenerative disorders and cancer (Herrando-Grabulosa et al., 2020; Kim & Maher, 2017; Maurice & Goguadze, 2017; Maurice & Su, 2009; Nguyen et al., 2017; D. A. Ryskamp et al., 2019). S1R agonists demonstrated neuroprotective effects in a variety of neurodegenerative disease models and are currently in clinical trials for a variety of neurological disorders including AD, HD, ALS, PD (Brimson, Brimson, Chomchoei, & Tencomnao, 2020; Herrando-Grabulosa et al., 2020; Maurice & Goguadze, 2017; Reilmann et al., 2019; D. A. Ryskamp et al., 2019). It has been proposed that accumulation of C99 fragment of APP leads to enhanced formation of cholesterol microdomains in the ER and upregulation of MAM activity in AD neurons (Area-Gomez et al., 2012; Montesinos et al., 2020; Pera et al., 2017). In contrast, MAM downregulation was observed in motor neurons in a genetic model of ALS (Watanabe et al., 2016). Our results suggest that S1R agonists allow remodeling of lipid microdomains in the ER membrane, which may help to normalize MAM function in AD and ALS neurons. Long-term effects of S1R activation may include more thorough remodeling of MAMs including size and protein composition needed for long-lasting metabolic adjustments in stress conditions. Changes in ER lipid microenvironment may also affect function of variety of channels, transporters and other signaling proteins localized to S1R signaling microdomains. The exact functional outcome of these lipid perturbations will be unique to each particular protein based on its activity in a lipid environment established by S1R.

In conclusion, we propose that many biological functions of S1R can be explained by its ability to organize and remodel cholesterol-enriched ER microdomains, which in turn affects activity of ER signaling proteins, stress-response and metabolic status of the cells.

## ACKNOWLEDGEMENTS

We are thankful to the members of Bezprozvanny laboratory for help and assistance with these studies and to Dr Arun Radhakrishnan for comments on the manuscript and useful suggestions. We are grateful to Drs Michael Hayden and Michal Geva (Prilenia Inc) for encouraging and supporting our work on S1R. JL is a Sowell Family Scholar in Medial Research. IB holds the Carl J. and Hortense M. Thomsen Chair in Alzheimer’s Disease Research. This work was supported by the National Institutes of Health R01NS056224 (IB), R01AG055577 (IB) and R01GM113079 (JL). MR is supported by the Howard Hughes Medical Institute and a grant from the Welch Foundation (I-1544).

## Author Contributions

V.Z.: Conceptualization; Data curation; Formal analysis; Investigation; Methodology, Writing – original draft
J.D.: Methodology, Writin – review & editing
W-R. L.: Investigation, Visualization
J.L.: Supervision, Resources, Funding Acquisition, Writing – review & editing
M.R: Supervision, Resources, Funding Acquisition, Writing – review & editing
l.B: Conceptualization, Supervision, Resources, Funding Acquisition, Writing – original draft, Writing – review & editing

## Declaration of Interests

The authors declare no competing interests.

## Figure supplements

**Figure 2 – figure supplement 1.**
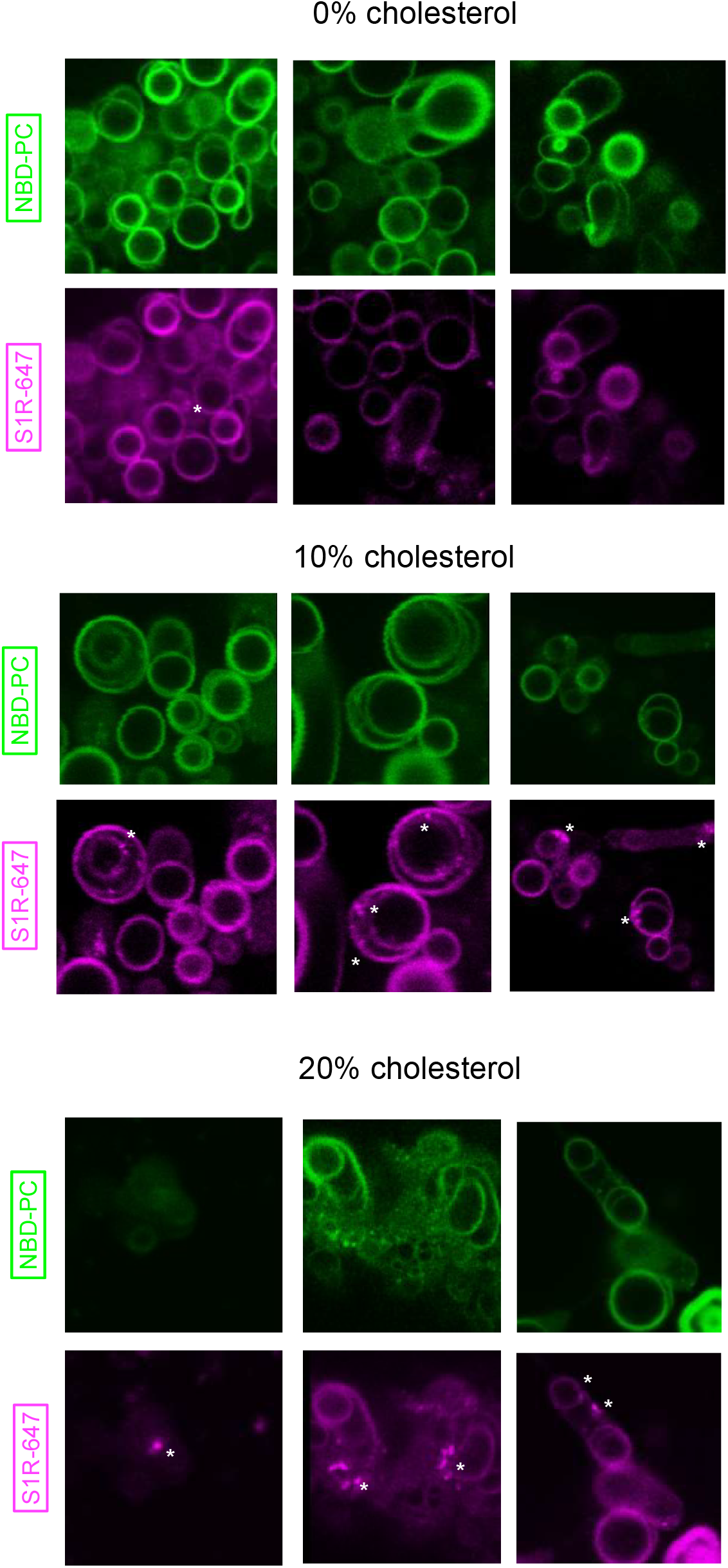
Additional examples of S1R distribution in DOPC GUVs at 0%, 10% and 20% mol cholesterol. Membrane dye NBD-PC is shown in green, S1R-647 is shown in magenta.

**Figure 3 – figure supplement 1.**
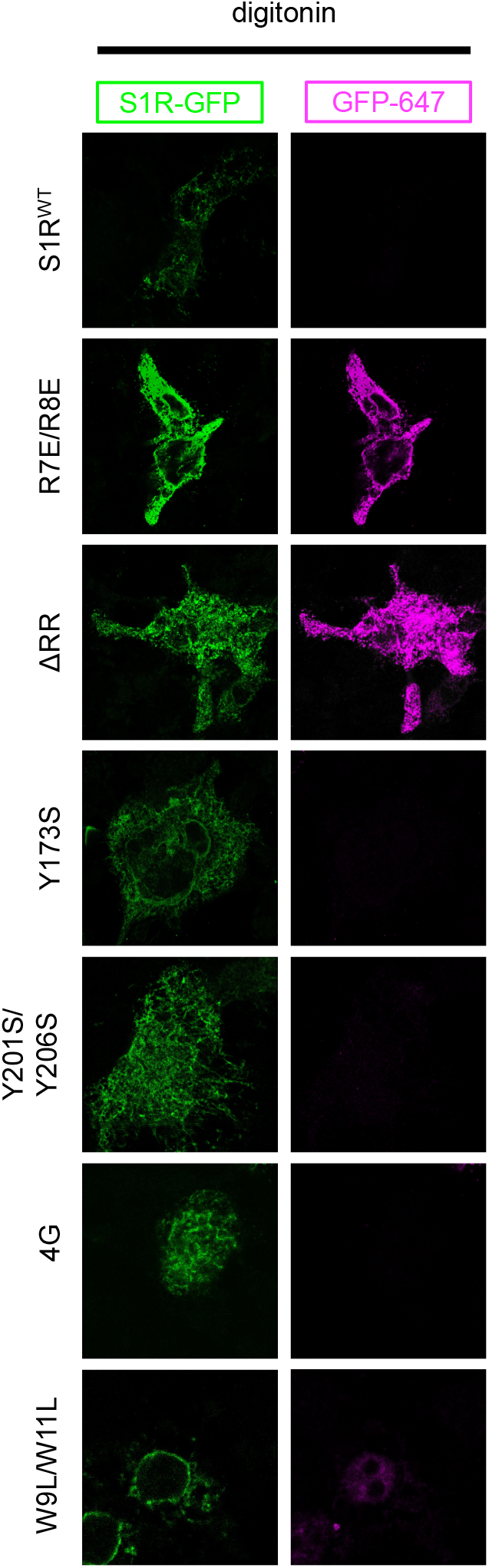
Membrane topologies of S1R-GFP proteins. HEK293 cells expressing WT S1R-GFP and mutants were permeabilized with digitonin (left) and stained with primary anti-GFP antibodies and Alexa647-conjugated secondary antibodies.

**Figure 4 – figure supplement 1.**
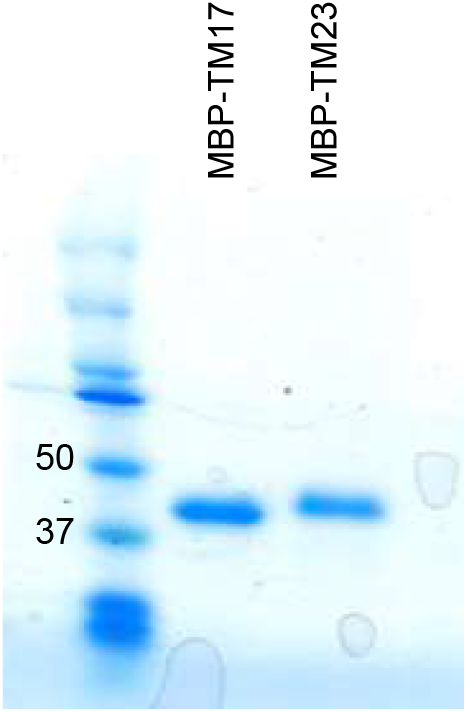
SDS-PAGE analysis of purified MBP-TM17 and MBP-TM27 proteins.

**Figure 5 – figure supplement 1.**
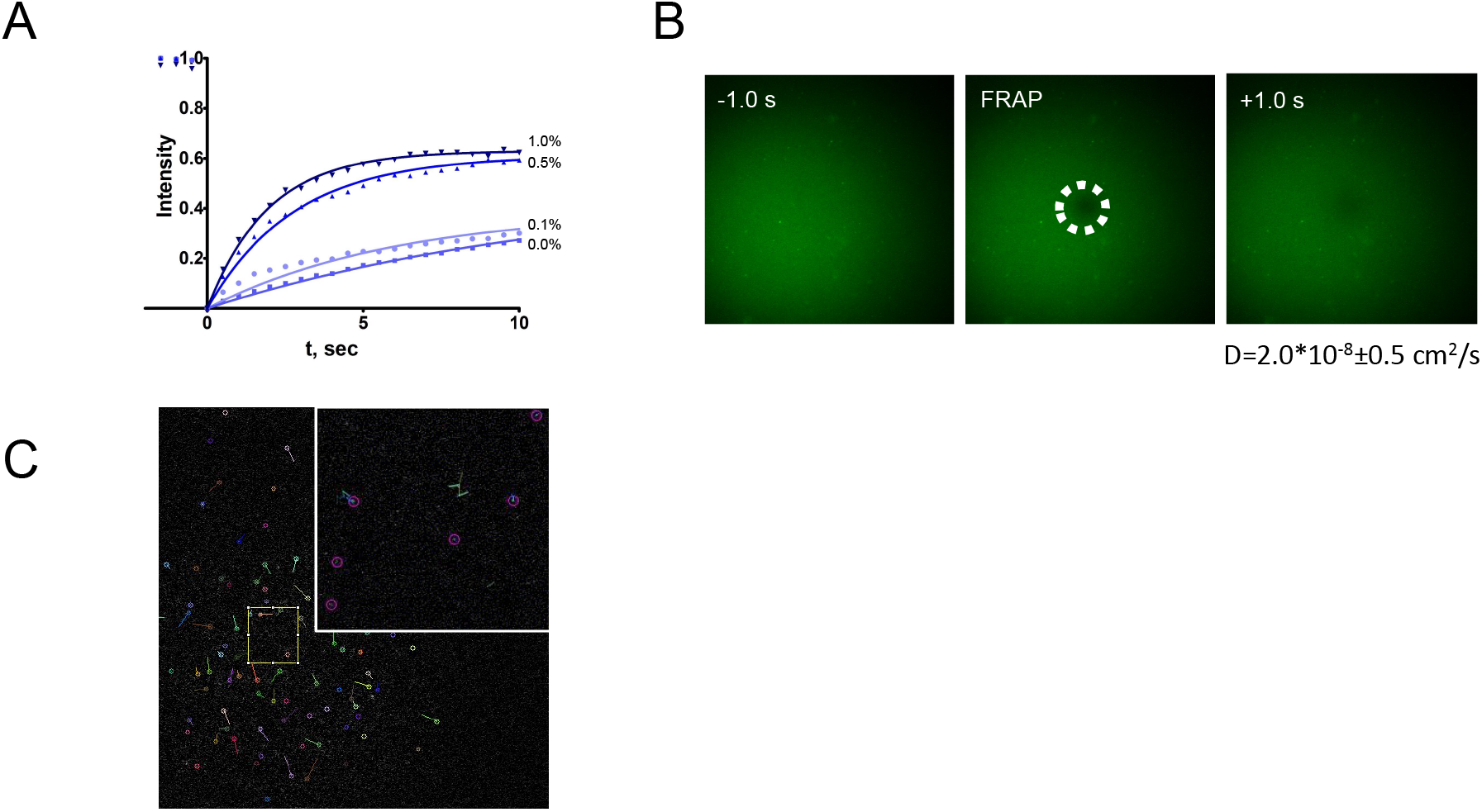
Characterization of DSLB technique. a. Lipid FRAP recovery curves of the second bilayer as a function of DSPE-PEG2000-biotin concentration. b. Recovery of the second bilayer after photobleaching, at 1.0% DSPE-PEG2000-biotin concentration c. Single-molecule detection of highly diluted S1R-647 at LPR 20,000:1.

**Figure 6 – figure supplement 1.**
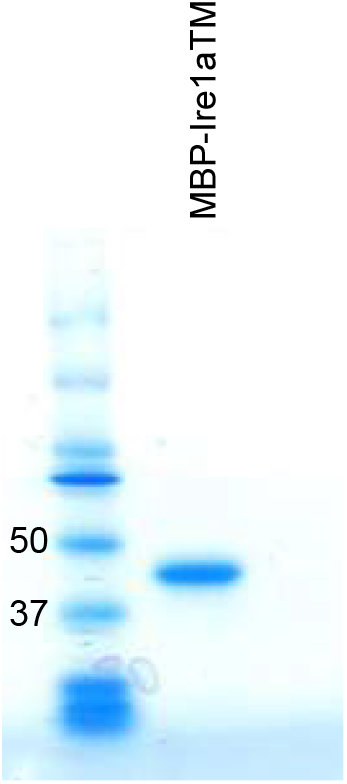
SDS-PAGE analysis of purified MBP-IRE1α-TM.

## MATERIALS AND METHODS

### Cell culture and transfection

HEK293T cells were cultured in DMEM medium supplemented with 10% fetal bovine serum. Transfection was performed using Lipofectamine LTX Plus according to manufacturer’s recommendations. HEK293T S1R-/-(S1R KO) cell line was generated in our previous work using CRISPR/Cas9-mediated deletion in the exon 1 of *SIGMAR1* gene (D. Ryskamp et al., 2017). Western blotting analysis confirmed efficient knockout of S1R in HEK293T cells.

### Lipids and detergents

Lipids: 1,2-dioleoyl-sn-glycero-3-phosphocholine (18:1 PC, DOPC), 1,2-dinervonoyl-sn-glycero-3-phosphocholine (24:1 PC, DNPC), 1-palmitoyl-2-{6-[(7-nitro-2-1,3-benzoxadiazol-4-yl)amino]hexanoyl}-sn-glycero-3-phosphocholine (NBD-PC), cholesterol, 1,2-distearoyl-sn-glycero-3-phosphoethanolamine-N-[biotinyl(polyethylene glycol)-2000 (DSPE-PEG(2000)-biotin), 1,2-dioleoyl-sn-glycero-3-[(N-(5-amino-1-carboxypentyl)iminodiacetic acid)succinyl] (nickel salt, DGS-NTA-Ni), L-a-phosphatidylcholine from chicken egg (Egg PC) were purchased from Avanti Polar Lipids. Detergents: n-dodecyl-N,N-dimethylamine-N-oxide (LDAO), n-dodecyl-β-D-maltopyranoside (DDM), n-octyl-β-D-glucopyranoside (OG), cholesteryl hemisuccinate (CHS) were purchased from Anatrace.

### Expression plasmids

Plasmid encoding human S1R gene fused with GFP (S1R-GFP) was generated by PCR amplification of human S1R gene (www.ncbi.nlm.nih.gov/nuccore/NM_005866.3) and cloning into pEGFP-N2 vector (Clontech) using HindIII/XbaI cloning sites. mCherry-Sec61β was obtained from Addgene (https://www.addgene.org/49155) (Zurek et al., 2011). Mutations were introduced using Q5 site-directed mutagenesis kit (NEB) according to manufacturer’s instructions. For baculovirus expression of 6*His-tagged S1R, human S1R gene was amplified by PCR and cloned into pFastBac-HTA vector (Bac-to-Bac baculovirus expression system, Thermo Fisher Scientific) using EcoRI/HindIII sites. Mutations were introduced using Q5 site-directed mutagenesis kit (NEB). Nucleic acid sequences encoding TM17, TM27 and human IRE1α-TM were synthesized by Genscript. Genes corresponding to the transmembrane peptides with following amino acid sequences: TM17 KKWWAAALLAAALLAAAWWKK and TM27 KKWWAAALLAAALLAAALLAAALLAAAWWKK, were synthesized by Genscript and cloned into pMAL-c5x vector using XmnI/NotI sites. A gene encoding a fragment of human IRE1α (a.a.r. 431-467 of the transmembrane helix and adjacent amphipathic helix), corresponding to the following amino acid sequence: ARPEAPVDSMLQDMATIILSTFLLIGWVAFIITYPLSK with a single point mutation K441Q, was synthesized by Genscript and cloned into pMAL-c5x using XmnI/NotI restriction sites. Genetic sequences of IRE1α, TM17 and TM27 were codon optimized for *E. coli* expression by Genscript. For cloning S1R-APEX2 fusion gene, APEX2 (https://www.addgene.org/49386) and human S1R genes were amplified by PCR. NotI site was introduced to the APEX2 5’ primer and to the S1R 3’ primer. PCR product was ligated using T4 ligase (NEB) and amplified using outer primers to produce the fusion gene S1R-APEX2. Resulting APEX2-S1R gene was cloned into lentivector expression plasmid (FUGW, addgene.org/14883).

Control plasmid encoding Sec61β-APEX2 (https://www.addgene.org/83411) was obtained from Addgene (S. Y. Lee et al., 2016) and APEX2-KDEL was generated by adding KDEL-encoding gene sequence to the 3’ reverse primer. Sec61β-APEX2 and APEX2-KDEL fusion genes were cloned into lentivector plasmid FUGW suing XbaI/EcoRI sites. All constructs were sequenced to confirm the accuracy of cloning.

### Protein expression, purification and NHS-conjugated dye labelling Purification of S1R

S1R was expressed with an N-terminal 6*His-tag fusion in Sf9 cells using Bac-to-Bac baculoviral expression system (Thermo Fisher Scientific) according to manufacturer’s recommendations. Infection was performed at 3*106 cells/ mL, cells were collected 68 h post-infection and cell pellet was stored at −80° C.

Cells were lysed in hypotonic buffer (20 mM HEPES pH=8.0, 1x cOmplete EDTA-free protease inhibitor cocktail (Roche) and sonicated three times for 2 minutes. Cell debris was centrifuged at 6,000 g for 15 minutes at 4° C. Supernatant was collected and pellet sonication was repeated one more time. After the second centrifugation step, supernatants were combined together and centrifuged at 100,000 g for 1 hour in a Ti70 rotor (Beckman Coulter). Membrane pellet was resuspended in a solubilization buffer (50 mM HEPES pH=8.0, 300 mM NaCl, 1% LDAO, 0.1% cholesterol hemisuccinate) using glass tissue homogenizer and rotated overnight at 4° C. Next day, solubilized membranes were centrifuged at 100,000 g for 1 hour. Supernatant was incubated with 1 mL of Ni2+-NTA agarose (Invitrogen) for 1 hour and washed with 25 mL of 50 mM HEPES pH=8.0, 300 mM NaCl, 1% LDAO, 0.1% cholesterol hemisuccinate, 10 mM imidazole and then with the same volume of buffer containing 20 mM imidazole. The protein was eluted in the same buffer containing 150 mM imidazole.

Receptor was further purified using anion-exchanged chromatography using HiTrap Q HP column (GE Healthcare). Protein was diluted 1:10 with buffer A (20 mM MOPS pH=7.0, 0.1% LDAO, 0.01% cholesterol hemisuccinate, 0.0015% Egg PC) and loaded onto a column, washed with 5 volumes of buffer A and eluted using linear gradient of buffer B (20 mM MOPS pH=7.0, 1 M NaCl, 0.1% LDAO, 0.01% cholesterol hemisuccinate, 0.0015% Egg PC). Fractions containing S1R were collected, pooled together and concentrated using Amicon centrifugal filters with 50,000 Da MWCO (Millipore).

Solution pH was adjusted to pH=8.0 with HEPES buffer and protein solution was mixed with an excess of Alexa647 NHS ester (Molecular probes) dissolved in DMSO, and left for labelling overnight at 4° C.

Next day, labelled protein was purified using gel-filtration chromatography on Superdex 200 10/300 column (GE Healthcare) in buffer containing 50 mM HEPES pH=8.0, 300 mM NaCl, 0.1% LDAO, 0.01% cholesterol hemisuccinate, 0.0015% Egg PC. Following the size-exclusion chromatography step, receptor was concentrated to 1-2 mg/mL, flash-frozen in small aliquots in liquid nitrogen and stored at −80° C.

### Purification of MBP-TM17, -TM27 and -IRE1α-TM

MBP-TM17, -TM27 and -IRE1α-TM were expressed in *E. coli* BL21(DE3) cells. 10 mL of LB medium were inoculated and grown overnight. Next day, the starter culture was added to 1 L of LB medium and grown until OD(600)=0.6. Protein synthesis was induced by addition of 0.3 mM IPTG. Proteins were expressed at 37° C for 3 h. Cells were collected by centrifugation and cell pellet was stored at −80° C.

Proteins were purified according to the procedure described in (Halbleib et al., 2017). Cell pellet was resuspended in lysis buffer (50 mM HEPES pH=7.4, 150 mM NaCl, 1 mM EDTA, 2 mM DTT, and 1x cOmplete protease inhibitor cocktail (Roche). Cells were sonicated three times for 2 minutes each. 500 mM OG stock was added to the final concentration of 50 mM and lysate was incubated for 10 minutes at 4° C. Lysate was clarified by centrifugation at 55,000 g for 1 h in 25.50 rotor (Beckman Coulter). Clarified supernatant was loaded on 1.5 mL of amylose resin (NEB) and washed with 50 mL of 50 mM HEPES pH=7.4, 150 mM NaCl, 1 mM EDTA, 2 mM DTT, 50 mM OG and then with 50 mL of buffer without DTT. Protein was eluted in the same buffer containing 10 mM maltose.

After concentration with Amicon centrifugal filters with 50,000 Da MWCO (Millipore), protein was labelled with Alexa555 NHS ester (Molecular Probes) as described for S1R. Protein was purified using Superdex 200 10/300 column (GE Healthcare) in a buffer containing 50 mM HEPES pH=7.4, 150 mM NaCl, 1mM EDTA, 0.018% DDM, concentrated, aliquoted, flash-frozen in liquid nitrogen and stored at −80° C.

### Purification of LAT

LAT was purified as described in (X. Su et al., 2016). BL21(DE3) cells containing MBP-His8-LAT 48-233-His6 were collected by centrifugation and lysed by cell disruption (Emulsiflex-C5, Avestin) in 20 mM imidazole (pH 8.0), 150 mM NaCl, 5 mM βME, 0.1% NP-40, 10% glycerol, 1 mM PMSF, 1 μg/ml antipain, 1 μg/ml pepstatin, and 1 μg/ml leupeptin. Centrifugation-cleared lysate was applied to Ni-NTA agarose (Qiagen), washed with 10 mM imidazole (pH 8.0), 150 mM NaCl, 5 mM βME, 0.01% NP-40, and 10% glycerol, and eluted with the same buffer containing 500 mM imidazole (pH 8.0). The MBP tag and His6 tag were removed using TEV protease treatment for 16 hrs at 4°C. Cleaved protein was applied to a Source 15 Q anion exchange column and eluted with a gradient of 200 mM-300 mM NaCl in 20 mM HEPES (pH 7.0) and 2 mM DTT followed by size exclusion chromatography using a Superdex 200 prepgrade column (GE Healthcare) in 25 mM HEPES (pH 7.5), 150 mM NaCl, 1 mM MgCl2, and 1 mM DTT. LAT was exchanged into buffer containing no reducing agent (25 mM HEPES (pH 7.0), 150 mM NaCl, 1 mM EDTA) using a HiTrap Desalting Column. C2-maleimide Alexa647 were added in excess and incubated with protein for 16 hrs at 4**°**C or 2 hrs at room temperature. Following the incubation, 5 mM 2-mercaptoethanol was added to the mixture to quench the labeling reaction. Excess dye was removed from labeled protein by size exclusion chromatography.

### Liposomal preparation and protein insertion for GUV preparations Reconstitution to liposomes

For protein reconstitution to liposomes, dry lipid films were prepared by dissolving lipids in chloroform with 0.1 mol % NBD-PC membrane label dye and drying overnight under vacuum.

Next day, films were rehydrated in 20 mM HEPES pH=8.0 buffer to the final concentration of 2.0 mg/mL and extruded using liposomal mini-extruder (Avanti Polar lipids) with 0.1 μm pore size polycarbonate filters (Avanti Polar Lipids). Liposomes were mixed with DDM (final 0.8 mM) and purified S1R at a lipid-to-protein ration of 200:1. Mixture was incubated at room temperature for 1 h. Detergent was removed by three addition of BioBeads SM-2 adsorbent resin (25 mg per 1 mL of lipid mixture) (BioRad) for 2 h each at 4**°**C. For DNPC reconstitution, initial incubation and BioBeads adsorbtion was performed at 27**°**C. Prepared liposomes were used on the day of experiment.

For co-reconstitution experiments with MBP-TM17, -TM27 and IRE1α-TM purified MBP fusion proteins were added at the initial incubation step at equimolar concentrations.

### Preparation of GUVs

GUVs were formed using polymer-assisted swelling on an agarose gel (Horger et al., 2009). Glass slides were covered with 1.0 % agarose and dried to completion on a hot plate. Small imaging chambers were assembled using adhesive silicon insulators (Electron Microscopy Sciences). Sucrose was added to the proteoliposomes at the final concentration of 15 mM, and 0.3 μL drops were deposited on the agarose-coated coverslips. After dehydration for 10 minutes at room temperature, slides were rehydrated in 50 mM HEPES pH=8.0, 150 mM NaCl buffer and incubated on a hot plate for 1 hour at 42**°**C. Samples were prepared side by side and imaged within 15 – 20 minutes after GUV formation. GUVs were imaged using upright fluorescent confocal microscope (Leica) with a 63x water immersion objective.

### Preparation of double supported lipid bilayers (DSLB)

For SUV preparation, lipid films of DOPC and 1.0 mol % DSPE-PEG(2000)-biotin were prepared and dried overnight under vacuum. Lipids were rehydrated in a bilayer buffer (50 mM Tris, pH=7.4, 150 mM NaCl, 1 mM TCEP) to the final concentration of 2.0 mg/mL and freeze-thawed ten times in liquid nitrogen. Liposomes were stored at −80°C. Prior to experiment, liposomes were centrifuged at 100,000 g for 1 h using a Sw55 rotor (Beckman Coulter), supernatant was collected and stored at 4**°**C up to two weeks under argon.

For the second bilayer mixtures, lipid films of DOPC, 1.0% mol DSPE-PEG(2000)-biotin, 0.1% NBD-PC and cholesterol at indicated molar concentrations were prepared and dried overnight under vacuum. Lipids were rehydrated to the final concentration of 2 mg/mL and extruded through the 0.1 μm polycarbonate filter as described above. Liposomes were mixed with mM DDM and purified sigma-1 receptor at lipid-to-protein ration 2000:1. Mixtures were incubated at room temperature for 1 hour and detergent was subsequently removed by three two-hour long additions of BioBeads SM-2 (25 mg per 1 mL of lipid mixture). Proteoliposomes were flash-frozen in liquid nitrogen and stored at −80 C. Prior to experiment, proteoliposomes were extruded through the 0.1 μm polycarbonate filter and centrifuged at 14,000 g for 10 minutes using a tabletop centrifuge.

Supported bilayers were formed in 96-well plates (MatriPlace MGB096-1-2LG-L, Brooks Life Science Systems). Glass surface was cleaned by immersing the plate in 5% Hellmanex III solution (Hellma Analytics) at 60**°**C for 3 hours. Plate was throughoutly washed with milliQ water to remove any remaining Hellmanex solution. Wells were dried with argon and sealed with foil tape. On the day of experiment, wells were cut open and hydrated with 500 μL milliQ water. 300 μL of 6 N NaOH were added to each well and plate was incubated on a heater for 1 h at 42**°**C. Then sodium hydroxide was removed and replaced with 300 μL of solution and incubated for one hour. After that, each well was washed three times with 700 μL of milliQ water and three times with 700 μL of bilayer buffer solution (50 mM Tris pH=7.4, 150 mM NaCl, 1 mM TCEP).

Each well was filled with 200 μL of the bilayer buffer and 20 μL of SUVs were added. Plate was incubated at 42**°**C for 2-3 hours and then each well was washed with 500 μL of the bilayer buffer. After that, proteoliposomes were added to each well (20 μL) and the plate was incubated at 42**°**C for 2 hours. Where indicated, ligands were added to the bilayer buffer during the second incubation step. Each well was washed ten times with 500 μL of the bilayer buffer.

For LAT-647 calibration experiments, bilayers were formed as described in (X. Su et al., 2016). Glass plates were cleaned as described above, washed and incubated with SUVs prepared by freeze-thaw method from 99% DOPC, 1% DGS-NTA-Ni and 0.1% NBD-PC. Bilayers were formed by described above. After washing three times with bilayer buffer, bilayer was blocked with 1 mg/mL BSA in bilayer buffer for 30 minutes at room temperature, washed three times and incubated with His-tagged LAT (final 0.1-1.0 pM) for 30 minutes and then washed to three times to remove unbound protein. TIRF images were captured using a Nikon Eclipse Ti microscope base equipped with an AndoriXon Ultra 897 EM-CCD camera with a 100 × 1.49 NA objective, a TIRF/iLAS2 TIRF/FRAP module (Biovision) mounted on a Leica DMI6000 microscope base equipped with a Hamamatsu ImagEMX2 EM-CCD camera with a 100 × 1.49 NA objective, or a Nikon Eclipse Ti microscope base equipped with a Hamamatsu ORCA Flash 4.0 camera with a 100 × 1.49 NA objective.

### Cell imaging

For imaging S1R localization in HEK293T cells, cells were cultured on glass coverslips in 24-well plates. Each well was transfected with 150 ng of S1R-GFP plasmid (WT or mutant) and 150 ng of mCherry-Sec61β using Lipofectamine LTX Plus (Thermo Fisher Scientific) according to manufacturer’s instructions. Cells were fixed 48 hours post-transfection in 4 % PFA/PBS solution for 20 minutes, permeabilized and blocked in 5% BSA, 0.1 % Triton X-100 in PBS for 1 hour and stained with anti-mCherry (16D7, 1:500, Invitrogen) and anti-TOM20 antibodies (FL-145, 1:500, Santa Cruz Biotechnology) overnight at 4**°**C. Next day, cells were washed and incubated with secondary antibodies (594 donkey anti-rat, A21209, 1:1,000, Invitrogen; 647 goat anti-rabbit, A27040, 1:1,000, Invitrogen) for 1 hour at room temperature. Cells were washed trice with PBS and mounted using Aqua Polymount solution (Polysciences). Cells were visualized using fluorescent confocal microscope (Leica) with 63x oil immersion objective.

### TIRFM imaging

HeLa cells were plated on 8-well Lab-Tek chambered coverglass (Nunc) at a density of 1.5 x 104 cells/well the day before transfection. Plasmid DNA were transfected into cells using TransIT-LT1 (Mirus Bio) with the 1:3 DNA-to reagent ratio. The plasmid DNA used in the transfection are: mTagBFP2-MAPPER (50ng/well) and GFP-S1R (50ng/well). Cells were washed with ECB before imaging and imaged in the ECB. TIRFM imaging experiments were performed at room temperature with a CFI Apo TIRF 100x/1.49 objective on a spinning-disc confocal TIRF microscope custom-built based on a Nikon Eclipse Ti-E inverted microscope (Nikon Instruments) with a HQ2 camera. The microscope was controlled by Micro-Manager software.

### Expansion microscopy

To prepare expanded specimens we used a procedure developed by Tillberg (Tillberg et al., 2016). Briefly, HEK293T cells were cultured, transfected with S1R-GFP plasmids and stained with primary (anti-GFP, ab13970, 1:500, Abcam and anti-TOM20, FL-145, 1:500, Santa Cruz Biotechnology) and secondary antibodies (488 goat anti-chicken, A11039, 1,1000, Invitrogen; Atto647N goat anti-rabbit 40839, 1:1,000, Sigma-Aldrich) as described above. Cells were processed according to protein-retention expansion microscopy (Tillberg et al., 2016). Briefly, permeabilized cells were incubated with 0.1 mg/mL 6-((acryloyl)amino)hexanoic acid, succinimidyl ester (AcX, Thermo Fisher Scientific) in PBS overnight, washed three times with PBS and incubated with gelation solution (1x PBS, 2 M NaCl, 8.625% (w/w) sodium acrylate, 2.5% (w/w) acrylamide, 0.15% (w/w) N,N′-methylenebisacrylamide, 0.02% TEMED, 0.02% ammonium persulfate (APS). Samples were transferred to 37**°**C tissue culture incubator for 2 hours. After gelation, samples were digested with proteinase K (NEB) diluted to 8 u/mL in digestion buffer (50 mM Tris pH=8.0, 1 mM EDTA, 0.5% Triton X-100, 1 M NaCl) overnight at room temperature. To enhance the fluorescent signal, samples were washed with PBS, blocked and re-stained with primary and secondary antibodies. Then, samples were transferred to milliQ water and allowed to complete expansion (about 1 hour with three water changes). Gel-embedded samples were mounted on poly-L-lysine-coated glass coverslips and visualized using fluorescent confocal microscope.

### APEX2 pull-down assay

For APEX2-based proximity-labelling experiments we followed a procedure described in (Hung et al., 2016). Briefly, HEK293T cells cultured on 10 cm2 dishes were transfected with 10 μg of S1R-APEX2, Sec61β-APEX2 or APEX2-KDEL plasmids. 48 hours post-transfection, cells were incubated in 500 μM biotin-phenol (Iris-Biotech) in complete medium at 37**°**C for 1 hour. Then, proteins were labeled by addition of 1 mM H_2_O_2_ for 1 minute and quenched with 10 mM sodium ascorbate, 5 mM Trolox, 10 mM sodium azide in PBS. Cells were lysed in RIPA buffer (50 mM Tris-HCl pH=7.4, 150 mM NaCl, 0.1% SDS, 0.5% sodium deoxycholate, 1% Triton X-100, 1x cOmplete protease inhibitor cocktail) for 15 minutes at 4 C on a rocker shaker. After centrifugation at 14,000 g for 10 minutes, 1 mL of lysate was mixed with 50 μL of streptavidin-agarose (Pierce) and incubated at 4**°**C for 4 hours on a rotary shaker. Resin was washed twice with 1 mL of RIPA buffer, once with 1 M KCl, once with 0.1 M Na2CO3, once with 2 M urea in 25 mM Tris-HCl pH=8.0, and twice with RIPA buffer. Proteins were eluted by boiling beads in 50 μL of 2x SDS Laemmli loading buffer plus 2 mM biotin. Eluted proteins were analyzed by Western blot.

### Western blot analysis

For UPR response experiments HEK293T cells were cultured in 6-well plates. Culture medium was replaced with complete medium and thapsigargin (Tg) (Calbiochem) was added at 1 μM concentration for the indicated periods of time. After that, culture medium was removed, and cells were processed for Western blot analysis. For Western blot analysis, cells were lysed in cold RIPA lysis buffer and incubated at 4**°**C for 10 minutes. Lysates were centrifuged at 14,000 g for 10 minutes, supernatants were collected and mixed with 6 x SDS loading buffer. Samples were boiled at 95**°**C for 10 minutes and proteins were separated by SDS-PAGE and analyzed by Western blotting with following antibodies: anti-phospho-IRE1α (NB100-2323, 1:10,000, Novus Biologicals), anti-IRE1α (14C10, 1:1,000, Cell Signaling), anti-XBP1s (E9V3E, 1:1,000, Cell Signaling), anti-tubulin (E7, 1:500, DSH), anti-APX2 (HRP) (ab192968, 1:1,000, Abcam), streptavidin-HRP (7403, 1:20,000, Abcam), anti-S1R (B-5, 1:300, Santa Cruz Biotechnology), anti-His tag (HIS.H8, 1:1,000, Millipore). The HRP-conjugated anti-mouse (111–035-144, 1:3,000) and anti-rabbit (115-035-146, 1:3,000) secondary antibodies were from Jackson ImmunoResearch.

### Quantification and statistical analyses

To calculate Mander’s colocalization coefficient between the ER (labeled with mCherry-Sec61β construct) and S1R-GFP proteins, JACOP plugin for ImageJ plugin was used (Bolte & Cordelieres, 2006). Acquisition parameters (such as laser power and gain) were kept constant for different samples. First, background subtraction was performed, and threshold was adjusted manually for S1R-GFP and mCherry-Sec61β channels. Thresholding was performed such as all the ER was segmented in the mCherry-Sec61β channel and S1R microdomains were clearly resolved in the S1R-GFP channel. For calculating mCherry-Sec61β:TOM20 and S1R-GFP:TOM20 colocalization in expanded samples Colocalization Highlighter plugin was used (Collins, 2007). Threshold levels were selected as mean signal intensities for mCherry-Sec61β and TOM20 channels, and manually adjusted for S1R-GFP channel to clearly segment microdomains (typically, at 1.5x mean intensity level). Colocalized areas were normalized to the total area occupied by mCherry-Sec61β or S1R-GFP at the same threshold level. Image data analysis of protein clustering in DSLB was performed using in-house MATLAB script. Fluorescent intensity of individual LAT-647 and S1R molecules were measured after background subtraction. Mean intensity of mono-cysteine-labelled LAT-647 molecules were used for calibration of the cluster size. For quantification of domain formation in double supported lipid bilayers, fluid bilayer areas were examined. Bilayer defects or unwashed liposomes were omitted from the analysis. To calculate the fraction of protein residing in clusters, we calculated integral intensity of S1R clusters in each field of view after background subtraction and divided it by the total protein density in each field of view. We considered clusters as protein assemblies with more that ten molecules of S1R. For Western blot analysis data were densitometrically analyzed using ImageJ software by normalizing the density of each band to IRE1α (for p-IRE1α quantification) or tubulin (for XBP1s quantification) of the same sample after background subtraction. The number of individual experiments, number of total cells or fields of view analyzed, and significance are reported in the figure legends. Statistical significance was calculated by Student’s t-test (for two-group comparison), one-way ANOVA with Tukey’s multiple comparison test (for comparison between different experimental groups) or Dunnett’s multiple comparison test (for comparison of multiple groups with one control group). The multiplicity adjusted p value is reported. p>0.05 = n.s, p<0.05 = *, p<0.01 = **, p<0.001 = *** and p<0.0001 = ****.

